# Revealing *In Situ* Molecular Profiles of Glomerular Cell Types and Substructures with Integrated Imaging Mass Spectrometry and Multiplexed Immunofluorescence Microscopy

**DOI:** 10.1101/2024.02.21.581450

**Authors:** Allison B. Esselman, Felipe A. Moser, Léonore Tideman, Lukasz G. Migas, Katerina V. Djambazova, Madeline E. Colley, Ellie L. Pingry, Nathan Heath Patterson, Melissa A. Farrow, Haichun Yang, Agnes B. Fogo, Mark de Caestecker, Raf Van de Plas, Jeffrey M. Spraggins

**Affiliations:** Mass Spectrometry Research Center, Vanderbilt University, Nashville, TN 37235, USA; Department of Chemistry, Vanderbilt University, Nashville, TN 37235, USA; Delft Center for Systems and Control, Delft University of Technology, 2628 CD Delft, The Netherlands; Department of Cell and Developmental Biology, Vanderbilt University, Nashville, TN 37232, USA; Department of Biochemistry, Vanderbilt University, Nashville, TN 37205, USA; Department of Pathology, Microbiology, and Immunology, Vanderbilt University Medical Center, Nashville, TN 37205, USA; Department of Pediatrics, Vanderbilt University Medical Center, Nashville, TN 37232, USA; Division of Nephrology and Hypertension, Department of Medicine, Vanderbilt University Medical Center, Nashville, TN 37232, USA

**Author notes:** Aspect Analytics, C-Mine 12, 3600 Genk, Belgium (Nathan Heath Patterson). Raf Van de Plas; Delft University of Technology, Delft Center for Systems and Control, Mechanical Engineering Faculty, Mekelweg 2 – Gebouw 34, 2628 CD, Delft, The Netherlands; M. Spraggins; 465 21^st^ Ave S. Room 9160, Medical Research Building III Vanderbilt University, Nashville, TN 37240.

**Keywords:** MALDI IMS, Immunofluorescence, Multimodal Imaging, Glomeruli, Cellular Analysis, Lipidomics

## Abstract

Glomeruli filter blood through the coordination of podocytes, mesangial cells, fenestrated endothelial cells, and the glomerular basement membrane. Cellular changes, such as podocyte loss, are associated with pathologies like diabetic kidney disease (DKD). However, little is known regarding the *in situ* molecular profiles of specific cell types and how these profiles change with disease. Matrix-assisted laser desorption/ionization imaging mass spectrometry (MALDI IMS) is well-suited for untargeted tissue mapping of a wide range of molecular classes. Additional imaging modalities can be integrated with MALDI IMS to associate these biomolecular distributions to specific cell types. Herein, we demonstrate an integrated workflow combining MALDI IMS and multiplexed immunofluorescence (MxIF) microscopy. High spatial resolution MALDI IMS (5 µm pixel size) was used to determine lipid distributions within human glomeruli, revealing intra-glomerular lipid heterogeneity. Mass spectrometric data were linked to specific glomerular cell types through new methods that enable MxIF microscopy to be performed on the same tissue section following MALDI IMS without sacrificing signal quality from either modality. A combination of machine-learning approaches was assembled, enabling cell-type segmentation and identification based on MxIF data followed by the mining of cell type or cluster-associated MALDI IMS signatures using classification models and interpretable machine learning. This allowed the automated discovery of spatially specific biomarker candidates for glomerular substructures and cell types. Overall, the work presented here establishes a toolbox for probing molecular signatures of glomerular cell types and substructures within tissue microenvironments and provides a framework that applies to other kidney tissue features and organ systems.

## Introduction

Kidney physiology is driven by highly organized multicellular functional tissue units (FTUs) that comprise the nephron. Probing the molecular profiles of specific tissue features across spatial scales, from entire FTUs to cell types, while maintaining spatial context, is critical for understanding both normal and diseased cellular functions. For example, glomeruli are complex FTUs that filter blood with the help of unique cell types, such as podocytes, mesangial cells, and fenestrated endothelial cells, organized around the glomerular basement membrane (GBM).^1^ Diseases of the kidney, such as diabetic kidney disease (DKD), can alter glomeruli, leading to podocyte loss, expansion of the mesangium, and thickening of the GBM.^2–4^ Determining the distribution of biomolecules among these cell types in healthy glomeruli is necessary to better understand cellular changes in diseased states.^5^ Imaging technologies are emerging to address the challenge of defining molecular characteristics of tissue features with increasing specificity.^6^ Several large-scale consortia, including the Human Biomolecular Atlas Program^7^ and the Kidney Precision Medicine Project^8^ are applying these approaches to construct molecular atlases of the human kidney.

Immunofluorescence microscopy is commonly used to map the distribution of specific proteins and delineate the cellular organization of tissues.^9^ Recent advancements in highly multiplexed methods allow more comprehensive spatial cell typing and the ability to reveal cellular organization. However, multiplexed immunofluorescence (MxIF) is targeted and limited to proteins, omitting other key molecular classes.^10^ Matrix-assisted laser desorption/ionization imaging mass spectrometry (MALDI IMS) addresses this limitation, by enabling untargeted, high spatial resolution imaging (<10 µm pixel sizes) of drugs, metabolites, lipids, glycans, and proteins, making it ideally suited for molecular discovery.^11,12^ Multimodal approaches combining MALDI IMS and MxIF allow hundreds of molecular features detected by IMS to be associated with specific tissue features and cell types.

MALDI IMS and MxIF are traditionally performed on serial sections. As modern instrumentation enables higher spatial resolution IMS, there is a greater need for multimodal imaging experiments to be performed on the same tissue section. While serial tissue sections remain appropriate for comparing larger-scale tissue features, cellular structures at a given location can change substantially between serial sections. Here, we demonstrate integrated methods for performing MALDI IMS and MxIF on a single tissue section, allowing IMS-reported molecular distributions to be directly correlated to MxIF-delineated tissue features, enabling automated discovery of *in situ* molecular marker candidates for glomerular cell types and substructures.

## Methods

Tissue sections were collected from a normal portion of a fresh-frozen renal cancer nephrectomy tissue and thaw-mounted onto glass slides. The sections were prepared for positive ion mode MALDI IMS, sampling only glomeruli as previously described.^13^ After IMS, sections were fixed for MxIF (cyclic-IF)^14^ analysis using 10 antibodies across 3 cycles. The immunofluorescence intensity signatures were clustered, segmenting the glomeruli into substructures, including specific glomerular cell types. Subsequently, a classification model was trained to recognize MxIF-based segments using IMS measurements as inputs. The model was interpreted using Shapley additive explanations (SHAP)^15,16^, and a global SHAP score was calculated for each of the IMS-measured molecular features, generating a ranked list of relevant biomarker candidates for each glomerular segment (i.e., cell type and substructure) See **Methods S1** for details.

## Results

High spatial resolution IMS methods were optimized to minimize tissue damage from laser irradiation. Following IMS acquisition, MxIF was performed to map specific cell types and structures within and surrounding the glomeruli (**Tables 1 and S1). Figures 1A** and **1B** show MxIF data from serial sections without and with a preceding IMS measurement. The comparison demonstrates the retention of MxIF stain quality post-MALDI IMS for the selected antibodies. Grayscale images of each antibody from both tissue sections are available in **Figures S1-S11**. IMS data were acquired first as the MxIF workflow can alter the molecular milieu of the tissue, reducing the capacity to perform subsequent spatial ‘omics experiments.

**Table 1.** Antibody Panel for MxIF Microscopy.

| Target | Cell Types/Structures | Cycle | Fluorophore | Conjugation Type |
| --- | --- | --- | --- | --- |
| Collagen IV $\alpha 1/2$ | Tubular Basement Membrane, Mesangial Matrix, and Glomerular Capsule | 1 | Cy5 | Indirect |
| Collagen IV $\alpha 5$ | Glomerular Basement Membrane, Glomerular Capsule, and Collecting Duct and Distal Convoluted Tubule Basement Membrane | 1 | Cy3 | Indirect |
| Tensin | Mesangial Cells and Vascular Smooth Muscle Cells | 1 | AF 488 | Indirect |
| Podocalyxin | Podocyte Cytoplasm and Plasma Membrane | 2 | AF 488 | Direct |
| Fibronectin | Mesangial Matrix and Muscularized Vessel Walls | 2 | AF 594 | Direct |
| CD31 | Endothelial Cells | 2 | AF 647 | Direct |
| Synaptopodin | Podocyte Cytoplasm and Plasma Membrane | 2 | Cy 7 | Direct |
| Nestin | Podocyte Cytoplasm | 3 | AF 488 | Direct |
| αSMA | Vascular Smooth Muscle Cells | 3 | AF 594 | Direct |
| AQP1 | Proximal Tubules | 3 | AF 647 | Direct |

**Figure 1.**
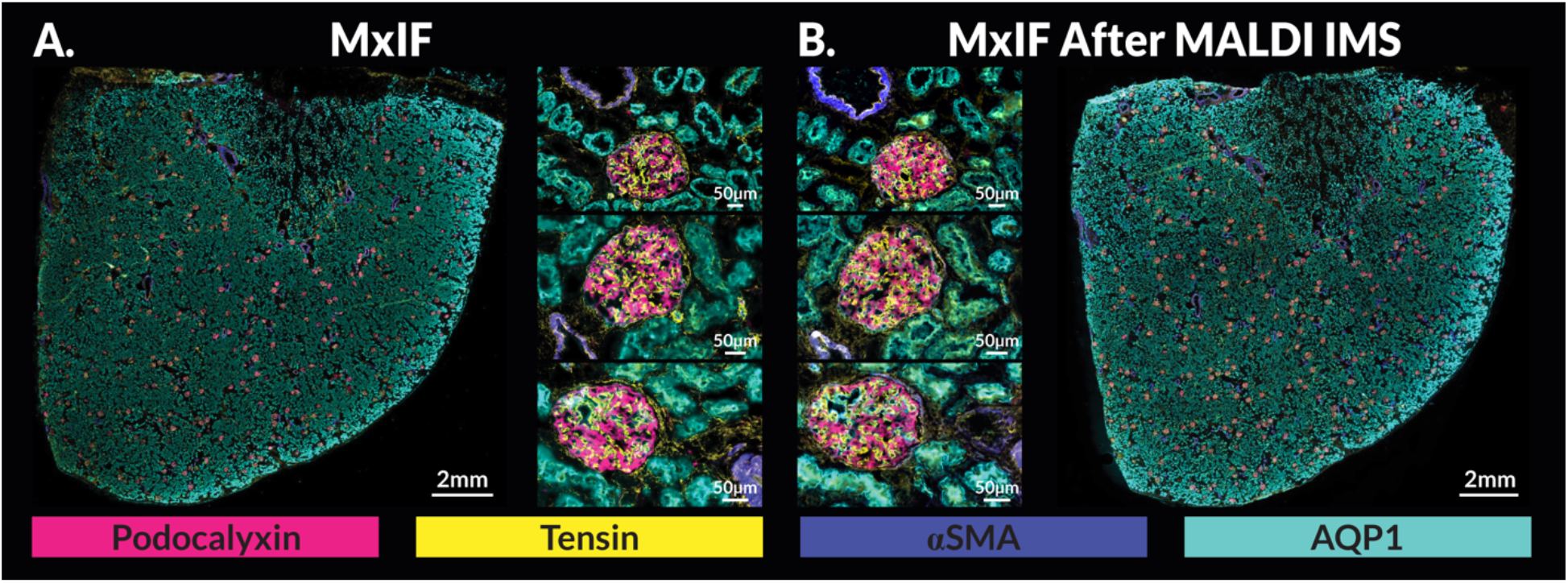
MxIF quality following MALDI imaging mass spectrometry. Comparison of serial human kidney tissue sections, one only stained and imaged using MxIF (**A**) and the other imaged with MALDI IMS targeting glomeruli and then stained and imaged using MxIF (**B**). Four of the ten antibodies are represented in the highlighted images. Selected regions from the whole slide images that include individual glomeruli (serial images of the same glomeruli) are provided to compare the stain quality between the two experiments. The stain quality of the four represented antibodies was minimally impacted by the MALDI IMS experiment and accompanying sample preparation. The 6 µm difference associated with the sectioning thickness between the serial sections shows differing patterns of podocalyxin (podocytes, pink) and tensin (mesangial cells, yellow) between the two serial sections, emphasizing the importance of performing multimodal imaging on the same tissue section.

Autofluorescence images were used to automatically define glomerular tissue areas for MALDI IMS measurement as described previously.^13,17^ This allowed for rapid acquisition of intra-glomerular lipid distributions for >250 glomeruli per tissue section.^13,17^ After IMS, tissue sections were stained with the antibody panel and imaged. **Figure S12** shows the whole-slide autofluorescence image, glomerular measurement regions, MALDI IMS, and MxIF of a single tissue section.

MxIF pixels within glomerular measurement regions were clustered using *k*-means based on the fluorescence intensities of tensin, podocalyxin, fibronectin, CD31, synaptopodin, and nestin, delineating 6 sub-glomerular segments that were enriched for specific glomerular cell types or substructures. The relationship between each segment and its corresponding cell type was based on its standardized mean antibody fluorescence intensity profile, as illustrated in **Figure S13**. Three glomeruli are highlighted in **Figure 2**, displaying lipid distributions uncovered by MALDI IMS (**Figure 2A)**, protein distributions from MxIF **(Figure 2B)**, and glomerular segments based on clustering of the MxIF data **(Figure 2C). Figure S14** provides data from additional example glomeruli from replicate samples, and **Figures S15-S26** display MxIF, glomerular segments based on clustering of MxIF, and MALDI IMS ion images for all glomeruli in each replicate. Data from all modalities were spatially co-registered, allowing the extraction of MALDI IMS pixels specific to each MxIF-based glomerular segment and the generation of an average mass spectrum for each associated substructure and cell type (**Figures S27-S32)**. The average differences in ion intensity between cluster segments (*i*.*e*., cell type or substructure) are shown in **Figure 3A-3B** and supplemental **Figures S33-S47**.

**Figure 2.**
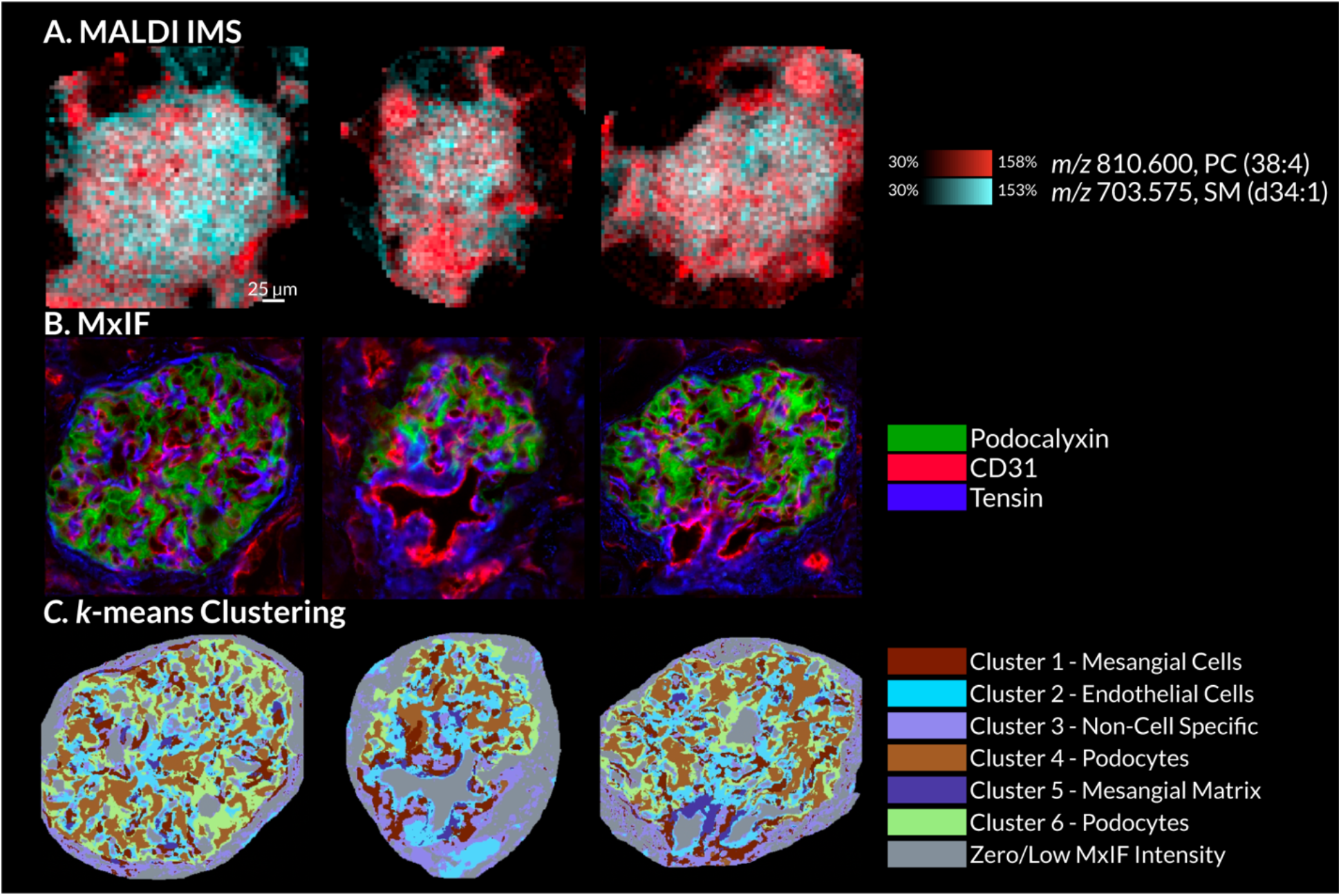
Multimodal molecular imaging data and segmentation maps from three selected glomeruli. MALDI ion images for PC(38:4) (*m/z* 810.600) and SM(d34:1) (*m/z* 703.575) highlight intra-glomerular molecular heterogeneity (**A**). Overlaid MxIF images of podocalyxin, CD31, and tensin (**B**). All *k*-means clustering-based segments of the MxIF data and the glomerular cell type or substructure that dominate each segment (**C**).

**Figure 3.**
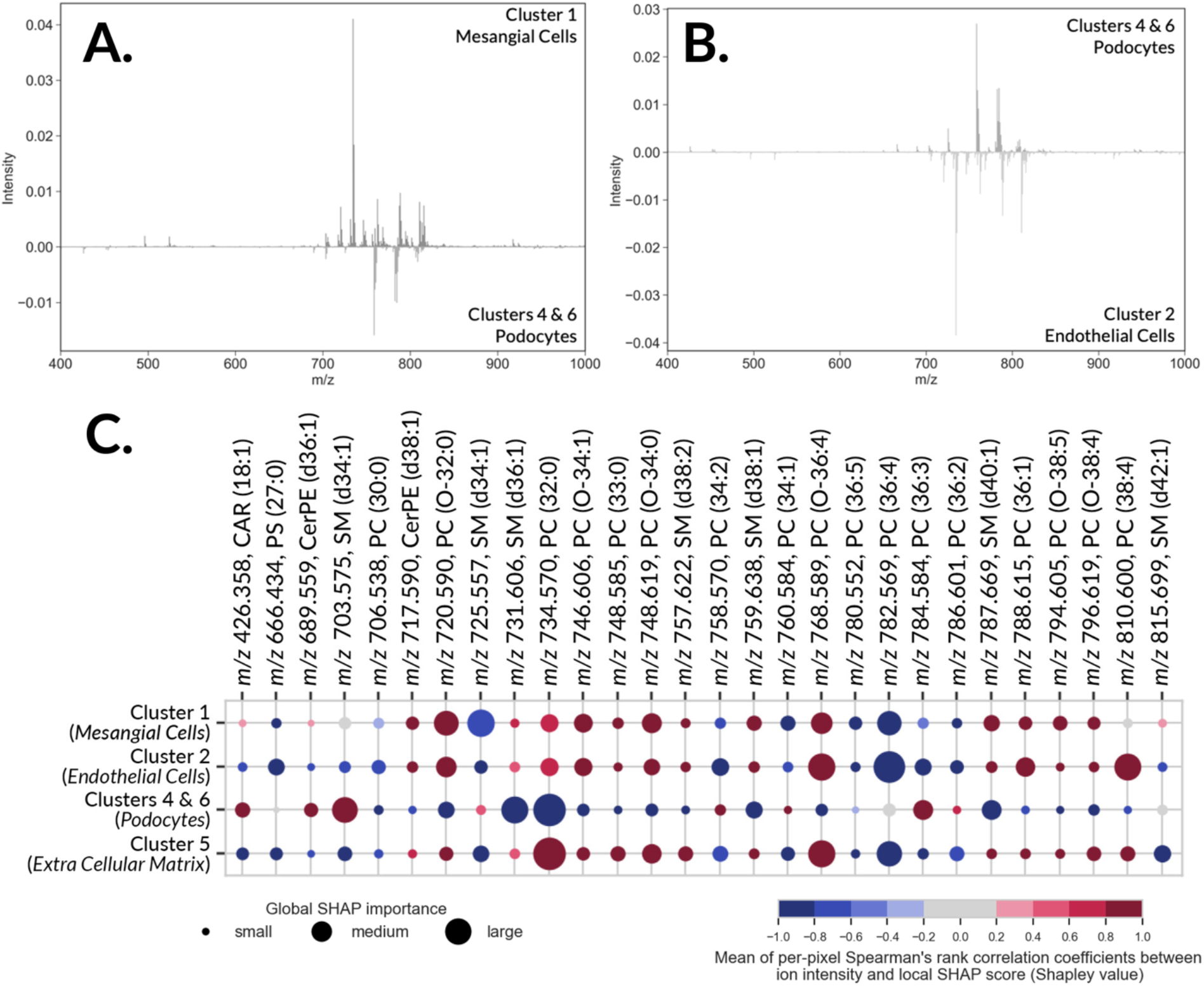
Molecular profiles specific to glomerular segments are revealed by integrating IMS and MxIF data. Mass spectral difference plots show the average ion intensity differences between cluster 1 (primarily mesangial cells) and combined clusters 4 and 6 (primarily podocytes) (**A**), and clusters 4 and 6 (primarily podocytes) versus cluster 2 (primarily endothelial cells) (**B**). The bubble plot from the SHAP analysis (**C**) summarizes the biomarker candidates for the glomerular segmentations and their dominant cell types and substructures. The size of each bubble indicates the global SHAP importance of a given ion species (column) to recognizing a given glomerular subarea (and its dominant cell type) (row), and the color indicates a positive (red) or negative (blue) correlation of the ion species abundance to that cluster’s recognition. From this analysis, it is shown that every glomerular segmentation has its own unique profile of relevant IMS-derived molecular species.

To discover multivariate molecular profiles distinctive for glomerular cell types and substructures, MALDI IMS and MxIF data were integrated using interpretable supervised machine learning. The MxIF-based segments were used as labels for each IMS pixel, and a classification model was trained to differentiate glomerular segments (and their dominant cell types) based on IMS-reported molecular ions. It is noted that the mean balanced accuracy, F1-score, precision, and recall were >85% for all segments (see **Table S5** for classification model performance metrics). Subsequently, SHAP was employed to interpret the model and discover biomarker candidates for each glomerular cluster.^15^ Given the model, SHAP ascertains the degree (relevance) and the direction (positive or negative correlation) of influence every ion has on the recognition of a particular glomerular segment.^15^ A global SHAP score was calculated for each IMS-reported molecule, quantifying its relevance to recognizing a certain segment, and providing a ranked list of biomarker candidates for each glomerular cell type or substructure (See **Figures S48-S51**). These data can be represented as a bubble plot (**Figure 3C**), where the size of each bubble indicates the global SHAP importance of a given molecule (column) for recognizing a given glomerular segment (row). The color indicates a positive (red) or negative (blue) correlation of the molecule’s abundance to the recognition of that segment. **Figure S52** shows the entire SHAP bubble plot, and **Table S6** summarizes the identification details for each molecule represented in the SHAP outputs.

## Discussion

We demonstrate an advanced multimodal workflow combining MALDI IMS, MxIF, and interpretable machine learning to uncover *in situ* molecular profiles of specific cell types and substructures of FTUs, here aimed at intra-glomerular features. Our methods maintain proper antibody staining post-MALDI IMS analysis, allowing the same tissue features to be sampled by multiple imaging modalities (not assured when using serial sections) and promoting conservative use of precious tissue samples. Multivariate SHAP analysis provides a unique fingerprint of positively and negatively correlated molecular species for every segmented tissue feature (**Figure 3C**). Interpreting the data in this way allows high-dimensional spatial ‘omics data to be mined efficiently and potential biomarker candidates to be readily discerned for each substructure or cell type. For instance, the phosphatidylcholine detected at *m/z* 810.600 (PC(38:4)) was determined to be a positively correlated biomarker candidate for the endothelial cell-related segment and a negatively correlated marker for the podocyte-related segment. Alternatively, sphingolipid SM(d34:1) (*m/z* 703.575) was a positively correlated biomarker candidate for the podocyte-related segment. Other sphingolipids have been shown to play a role in podocyte homeostasis, mediating normal and disease-related responses.^18^ The ceramide chain of SM(d34:1) is synthesized by CERS6. CERS6 expression is critical for podocyte cytoskeletal organization and maintenance of slit diaphragms.^19^ Based on our observations of SM(d34:1) as a robust glomerular biomarker coupled with the previously known molecular relationship of CERS6 with podocytes, we hypothesize that SM(d34:1) is a critical lipid regulator of podocyte structure and function. The ability to unveil these connections between biomarkers and known biology points to the potential for our integrated approach to serve as a molecular discovery tool for advancing the understanding of mechanisms of cellular function within tissue microenvironments.

As the most complex kidney FTU, glomeruli serve as a challenging case study demonstrating the broad applicability of our workflow. It can be readily adapted to analyze other FTUs, including tubules and ducts in the kidney, as well as other organ types. While this proof-of-concept study was performed on healthy tissue, it could be applied to study diseases such as chronic kidney disease. For example, as DKD progresses, glomeruli undergo podocyte loss and mesangial cell expansion, but the cellular-level lipidomic changes that occur are not well characterized.^20^ Our workflow can be used to find spatially specific biomarker candidates for renal cell types, which could be used to subtype DKD and other kidney diseases. In essence, our toolbox offers insight into the complex relationship between the molecular and cellular organization of tissues, paving the way for precision medicine by uncovering how these relationships are altered in normal aging and disease.

## Acknowledgments

This work was supported by the National Institutes of Health (NIH) Common Fund and National Institute Of Diabetes And Digestive And Kidney Diseases (NIDDK) under Award Numbers U54DK134302 and U01DK133766 (J.M.S. and R.V.), by the NIH Common Fund and National Eye Institute (NEI) under Award Number U54EY032442 (J.M.S. and R.V.), by the NIH National Institute On Aging (NIA) under Award Number R01AG078803 (J.M.S. and R.V.), and by the National Science Foundation Major Research Instrument Program CBET – 1828299 (J.M.S.). This research was furthermore made possible by the Chan Zuckerberg Initiative DAF, an advised fund of Silicon Valley Community Foundation, under Award Numbers 2021-240339 and 2022-309518 (L.G.M. and R.V.). K.V.D was supported by an NIDDK training grant (T32DK007569-34). The content is solely the responsibility of the authors and does not necessarily represent the official views of the funders. The authors would like to thank Jamie Allen for processing and preparing frozen tissue blocks of human kidney tissue.

## Notes

### Competing Interest Statement

The authors have declared no competing interest.

