## Supplementary material for "Revealing *In Situ* Molecular Profiles of Glomerular Cell Types and Substructures with Integrated Imaging Mass Spectrometry and Multiplexed Immunofluorescence Microscopy": Methods S1

[#]Current affiliation: Aspect Analytics, C-Mine 12, 3600 Genk, Belgium

### Table of Contents


| | | |
|---|---|---|
| Method S1 | Extended Methods | S-3 – S-8 |
| Table S1 | Antibody Information | S-9 |
| Table S2 | qTOF Instrument Parameters | S-10 |
| Table S3 | LC-MS/MS Negative Ion Mode Instrument Parameters | S-10 |
| Table S4 | LC-MS/MS Positive Ion Mode Instrument Parameters | S-11 |
| Figure S1 | Grayscale Antibody Images of DAPI | S-11 |
| Figure S2 | Grayscale Antibody Images of Collagen IV α5 | S-12 |
| Figure S3 | Grayscale Antibody Images of Collagen IV α1/2 | S-12 |
| Figure S4 | Grayscale Antibody Images of Tensin | S-12 |
| Figure S5 | Grayscale Antibody Images of Podocalyxin | S-13 |
| Figure S6 | Grayscale Antibody Images of Fibronectin | S-13 |
| Figure S7 | Grayscale Antibody Images of CD31 | S-13 |
| Figure S8 | Grayscale Antibody Images of Synaptopodin | S-14 |
| Figure S9 | Grayscale Antibody Images of Nestin | S-14 |

| | | |
|---|---|---|
| Figure S10 | Grayscale Antibody Images of αSMA | S-14 |
| Figure S11 | Grayscale Antibody Images of AQP1 | S-15 |
| Figure S12 | Whole Slide Imaging Workflow | S-15 |
| Figure S13 | Standardized Mean Fluorescent Intensity Bar Plot | S-16 |
| Figure S14 | Example Glomeruli from Three Serial Sections | S-16 |
| Figure S15 | MxIF Mosaic of Glomeruli From Section 1 | S-17 |
| Figure S16 | *k*-means Clustering Mosaic of Glomeruli From Section 1 | S-18 |
| Figure S17 | SM (d34:1) Ion Image Mosaic of Glomeruli From Section 1 | S-19 |
| Figure S18 | PC (38:4) Ion Image Mosaic of Glomeruli From Section 1 | S-20 |
| Figure S19 | MxIF Mosaic of Glomeruli From Section 2 | S-21 |
| Figure S20 | *k*-means Clustering Mosaic of Glomeruli From Section 2 | S-22 |
| Figure S21 | SM (d34:1) Ion Image Mosaic of Glomeruli From Section 2 | S-23 |
| Figure S22 | PC (38:4) Ion Image Mosaic of Glomeruli From Section 2 | S-24 |
| Figure S23 | MxIF Mosaic of Glomeruli From Section 3 | S-25 |
| Figure S24 | *k*-means Clustering Mosaic of Glomeruli From Section 3 | S-26 |
| Figure S25 | SM (d34:1) Ion Image Mosaic of Glomeruli From Section 3 | S-27 |
| Figure S26 | PC (38:4) Ion Image Mosaic of Glomeruli From Section 3 | S-28 |
| Figure S27 | Average Mass Spectrum of Cluster 1 | S-29 |
| Figure S28 | Average Mass Spectrum of Cluster 2 | S-29 |
| Figure S29 | Average Mass Spectrum of Cluster 3 | S-30 |
| Figure S30 | Average Mass Spectrum of Clusters 4 and 6 | S-30 |
| Figure S31 | Average Mass Spectrum of Cluster 5 | S-31 |
| Figure S32 | Average Mass Spectrum of Cluster 0 | S-31 |
| Figure S33 | Plot of Spectral Differences between Cluster 1 and 2 | S-32 |
| Figure S34 | Plot of Spectral Differences between Cluster 1 and Clusters 4 and 6 | S-32 |
| Figure S35 | Plot of Spectral Differences between Cluster 3 and 1 | S-33 |
| Figure S36 | Plot of Spectral Differences between Cluster 3 and 2 | S-33 |
| Figure S37 | Plot of Spectral Differences between Cluster 3 and Clusters 4 and 6 | S-34 |
| Figure S38 | Plot of Spectral Differences between Cluster 3 and 5 | S-34 |
| Figure S39 | Plot of Spectral Differences between Cluster 3 and 0 | S-35 |
| Figure S40 | Plot of Spectral Differences between Clusters 4 and 6 and Cluster 2 | S-35 |
| Figure S41 | Plot of Spectral Differences between Cluster 5 and 1 | S-36 |
| Figure S42 | Plot of Spectral Differences between Cluster 5 and 2 | S-36 |
| Figure S43 | Plot of Spectral Differences between Cluster 5 and Clusters 4 and 6 | S-37 |
| Figure S44 | Plot of Spectral Differences between Cluster 5 and 0 | S-37 |
| Figure S45 | Plot of Spectral Differences between Cluster 0 and 1 | S-38 |
| Figure S46 | Plot of Spectral Differences between Cluster 0 and 2 | S-38 |
| Figure S47 | Plot of Spectral Differences between Cluster 0 and Clusters 4 and 6 | S-39 |
| Table S5 | SHAP Performance Metrics | S-39 |
| Figure S48 | Ranked SHAP Values for Cluster 1 | S-40 |
| Figure S49 | Ranked SHAP Values for Cluster 2 | S-40 |
| Figure S50 | Ranked SHAP Values for Clusters 4 and 6 | S-40 |
| Figure S51 | Ranked SHAP Values for Cluster 5 | S-41 |
| Figure S52 | Full, Unlabeled SHAP Bubble Plot | S-41 |
| Table S6 | Lipid Identification Table | S-42 |

**Method S1**

**<u>Materials</u>.** Ammonium formate, carboxymethylcellulose sodium (CMC), glycerol, 30% hydrogen peroxide, sodium hydroxide, 10% neutral buffered formalin (NBF), and a periodic acid-Schiff (PAS) stain kit was purchased from Sigma-Aldrich (St. Louis, MO). The matrix 4-(dimethylamino)cinnamic acid (DMACA) with 99% purity and 20mM Hoechst 3342 were bought from Thermo Scientific (Waltham, MA). High-performance liquid chromatography (HPLC)-grade acetone, isopentane, and Superfrost Plus microscope slides were purchased from Fisher Scientific (Pittsburg, PA). Universal blocking reagent (10x) was purchased from BioGenex Laboratories (Fremont, CA), and antibody diluent reagent solution was purchased from Invitrogen (Waltham, MA). Secondary antibodies: anti-rat conjugated with Cy3, anti-mouse conjugated with Cy5, and anti-rabbit conjugated with AF 488 were purchased from Jackson Immuno Research (West Grove, PA). A Lightning-Link antibody conjugation kit for Cy7 was purchased from Abcam (Cambridge, UK). Equisplash (330731), cardiolipin (791108C), and sulfatide (860736) standards were purchased from Avanti Polar Lipids (Birmingham, AL). Water, methanol, acetonitrile, 2-propanol, methyl tert-butyl ether (MTBE), and ammonium formate (for LC-MS/MS) were purchased from EMD Millipore (Burlington, MA); formic acid (for LC-MS/MS) was purchased from Thermo Scientific (Rockford, IL). Human kidney samples were provided by the Cooperative Human Tissue Network at Vanderbilt University Medical Center. **Table S1** provides additional antibody information.

**<u>Sample and Data Collection</u>.** De-identified human kidney samples were obtained from a disease-free tumor associated nephrectomy from a 57-year-old white male with a BMI of 25.3. Samples were collected by the Vanderbilt Cooperative Human Tissue Network (CHTN, IRB Protocols #181822 "Biomolecular Multimodal Imaging for 3-Dimensional Tissue Mapping of the Human Kidney") and placed on ice within 1-2 hours of surgery. Tissue blocks (~0.5cm × 1cm × 2-4cm) were cut extending from the cortex through the medulla on their long axis, and frozen in 2.6% CMC in water on a dry ice/isopentane slurry and stored at -80C prior to sectioning.[1] Serial 6 μm thick cryosections were cut using a CM3050 S cryostat (Leica Biosystems, Wetzlar, Germany) and thaw-mounted onto Superfrost Plus microscope slides. Autofluorescence images were acquired of each section using standard DAPI, eGFP, and DSRed fluorescent filters on a Zeiss

AxioScan.Z1 slide scanner (Carl Zeiss Microscopy GmbH, Oberkochen, Germany), equipped with a Colibri7 LED light source.

**<u>MALDI IMS Preparation</u>.** Three serial sections were prepared for MALDI IMS analysis. The sections were washed with chilled (4 °C) 150 mM ammonium formate 3 times for 45 seconds each, then dried with nitrogen gas. An in-house developed sublimation device was used to sublime 5 mg of 4-(dimethylamino)cinnamic acid (DMACA) onto the slide. The apparatus was heated (190 °C) for 10 minutes under vacuum (110-150 mTorr) while the sample was cooled to -78 °C using a dry ice and acetone slurry in the cold finger, resulting in a final matrix density of ~0.22 µg/mm$^2$.[2] Functional tissue unit (FTU) segmentation of glomeruli was performed using the autofluorescence image of each tissue section using a previously described method.[3–5] The FTU segmentations were used to define measurement regions so that only the glomeruli were measured by MALDI IMS. The FTU segmented areas were scaled by a factor 1.6 using an affine scale transform to allow for slight errors in spatial targeting and to ensure capture of the border of each glomerulus. The MALDI IMS experiments were performed on a timsTOF fleX mass spectrometer equipped with a microGRID stage (Bruker Daltonik, Bremen, Germany).[6] Tissue imaging data were collected at 5 µm × 5 µm pixel size with no beam scan, a 5 µm pitch, 25 shots per pixel, and 4.8% total laser power. Lipid data were collected in positive ion mode from *m/z* 400 to 2000. Instrument-specific parameters are available in **Table S2**. Molecular annotations were supported by LC-MS/MS. Selected IMS peaks were linked to LC-MS/MS data using mass accuracy (<5 ppm). After MALDI IMS data acquisition, post-IMS autofluorescence images were acquired on the same tissue sections prior to matrix removal using a Zeiss AxioScan.Z1 fluorescence slide scanner using the previously described eGFP fluorescence filter and a monochromatic brightfield image. After the collection of the post-IMS autofluorescence images, all tissue sections underwent the MxIF protocol.

**<u>LC-MS/MS</u>.** Two fresh frozen 10 µm thick tissue sections were added to glass vials along with 5 µL of 100 µg/mL internal standards Avanti Equisplash, C24:1 mono-sulfo galactosyl(ß) ceramide-d7 (d18:1/24:1) (Avanti), and 18:2 Cardiolipin-d5 (Avanti) and methanol was added to cover the tissue. Metal beads were added to the vials and the tissue was homogenized by vortexing for 1 minute, followed by 5 minutes on dry ice and 30 minutes sonication on ice. Lipids were extracted by adding 1600 µL of cold MTBE and 400 µL of cold water to 400 µL of cold methanol. Vials were centrifuged at 100 × *g* for 10 minutes at 4 °C and then allowed to rest on ice for 10 minutes. The top layer (1400 μL) was extracted to a new vial and dried under nitrogen. The sample was resuspended in 300 μL of methanol. Reversed-phase separation was performed on a Waters Premier UHPLC with a 2.1×100 mm Waters Premier CSH-C18 column heated to 60 °C. Mobile phase A consisted of 60:40 ACN:$H_2O$ with 10 mM ammonium formate and 0.1% formic acid. Mobile Phase B consisted of 90:10 IPA:ACN with 10 mM ammonium formate and 0.1% formic acid. The total analysis time was 20 minutes, followed by a 10 minute equilibration time. A timsTOF fleX mass spectrometer utilized parallel accumulation-serial fragmentation (PASEF) and MS/MS stepping to obtain the collision cross section (CCS) of each species to permit the simultaneous measurement of both low *m/z* fragments and higher molecular weight lipids. Instrument-specific parameters can be found in **Table S3** and **Table S4.** Analysis was performed with MS-DIAL version 4.80 and all lipid annotations were blank filtered, retention scored, and referenced matched with the MS-DIAL combined database.

**Cyclic MxIF and Histological Stain Preparation.** Cyclic MxIF was performed using a modified protocol.[7] Briefly, tissue sections were fixed with 10% NBF for 5 minutes. The tissue sections were then washed with 1X PBS, blocked for 30 minutes with 1X universal blocking reagent, and then incubated with the first cycle of antibodies overnight at 4 °C (see **Table S1** for MxIF cycle and antibody information). Tissue sections were then washed with 1X PBS and incubated with secondary antibodies for 60 minutes for any unconjugated primary antibodies. The sections were washed with 1X PBS and incubated with Hoechst for 10 minutes before attaching a coverslip using a 50:50 glycerol:$H_2O$ solution. Each cycle was imaged using a Zeiss AxioScan.Z1 fluorescence slide scanner using standard AF488, AF594, AF647, Cy3, Cy5, Cy7, and DAPI filters. The coverslips were removed by soaking the slides in 1X PBS. The fluorophores were inactivated between each cycle by laying the slides over an LED light and pipetting a solution of basic 0.1 M sodium bicarbonate with 3% hydrogen peroxide in deionized water. After an hour, the slides were washed with 1X PBS and incubated overnight at 4 °C with the next cycle of antibodies. This process continued until 3 cycles had been imaged. Histograms from each image were adjusted individually as best fit for **Figures S1-S11**. After the last cycle was imaged, the coverslip was removed with 1X PBS, and the tissue section was stained with PAS.[8]

**Multimodal Image Registration and Data Analysis.** MALDI IMS and microscopy images were registered in a two-step process. First, MALDI IMS pixels were aligned to laser ablation marks, as measured by the post-IMS autofluorescence image, using in-house developed software *image2image* to manually select corresponding pairs of laser ablation marks and IMS pixels. Additional microscopy images were then registered to the IMS data via the previously registered post-IMS autofluorescence images in the *wsireg* software.[9] All registered whole-slide images were stored in the vendor-neutral pyramidal OME-TIFF format and maintained their original spatial resolution (i.e., there is no downsampling or loss of pixel spacing through the registration process). After alignment of all images, automated glomeruli segmentations, as predicted by a convolutional neural network, were used to find the MALDI IMS pixels associated with each detected glomerulus and were extracted for further data analysis.

***k*-means Clustering of Glomeruli Pixels.** Cell types were segmented by clustering the fluorescent intensities of MxIF antibodies. First, glomeruli specific antibody channels (tensin, podocalyxin, fibronectin, CD31, synaptopodin, and nestin) of each image were manually thresholded to remove the majority of the background and non-specific binding intensities. Then, the previously described glomeruli segmentations were used to mask the glomeruli of each image, resulting in a total of 67,269,568 pixels of N=6 dimensions (one for each of the channels above). The 27,491,379 pixels that were below the manual threshold across all channels were separated into their own cluster. The intensities of the remaining pixels were standardized on a per-channel basis before clustering them using a standard *k*-means algorithm. This was repeated for clusters ranging from *k*=2 to *k*=15, the optimal of which was determined to be *k*=6 by using the elbow method. The segmentation masks were then generated by placing a different integer value between 1 and 6 in the location of each pixel in the original image, depending on which cluster they belong to. A value of 0 was given to the thresholded pixels that were excluded from the *k*-means clustering to distinguish them from the rest. Finally, each class of the segmentation masks was refined by performing a single iteration of binary dilation followed by a single iteration of binary erosion and removing objects that were smaller than 6 pixels.

**MALDI IMS Data Preprocessing.** MALDI IMS data were exported from Bruker timsTOF file format (.d) to a custom binary format. Each pixel/frame contained centroid peaks that covered the entire acquisition range and was reconstructed to form a pseudo-profile mass spectrum using Bruker's SDK (v2.21). The data were *m/z* aligned using 6 peaks that appeared in at least 50% of the pixels using the *msalign* library (v0.2.0). The mass axis of the dataset was calibrated, using theoretical masses for the 6 peaks, to approximately ±1 ppm precision. The MALDI IMS data was subsequently normalized using a total ion current (TIC) approach. Then, an average mass spectrum based on all pixels belonging to each *k*-means cluster in the data set was computed. The average mass spectrum was peak-picked and a total of 1161 peaks were detected. Note that isotopic peaks were not removed in the classification workflow.

**Supervised Machine Learning and Shapley Additive Explanations.** Clusters 1, 2, 3, 5, and the combination of 4 and 6 were used as classes to build eXtreme Gradient Boosting (XGBoost) classification models that recognize one of the 5 clusters. We take the one-versus-all approach to multiclass classification: each classification task consists of differentiating one cluster from the other four clusters (imbalanced binary classification). Our chosen interpretability method, Shapley additive explanations (SHAP), enables us to determine which ion species have a marker-like relationship to each cluster by quantifying each ion species' importance to the corresponding classification models and the task of recognizing a certain MxIF-delivered glomerular cluster. For a given classification (cluster recognition) task, SHAP measures the global (experiment-wide) and local (per-pixel) relevance, or importance, of each ion species. The global SHAP importance scores (hereafter referred to as SHAP scores) are used to rank all IMS-provided ion species by decreasing relevance to recognizing a specific cluster, and the top of the list can be used to select a subset of highly discriminative molecular species that represent potential biologically significant marker candidates for the cluster in question. The local SHAP importance scores, or local Shapley scores, are used to measure the direction of relevance (by positive or negative monotonic correlation) and to assess the significance of the relationship between a given molecular species' ion intensity measurements and a pixel's likelihood of belonging to a given cluster.

**Classification.** The first step of the workflow is cluster recognition. Rather than use one XGBoost model, we use an ensemble of 10 XGBoost models for each one-versus-all classification task. The training process of each of these XGBoost models is initialized using a different random seed, and the models are trained on slightly different training datasets. The ten training datasets differ in the following two ways: the random under-sampling of the majority class (and, in case of extreme class imbalance, the oversampling of the minority class by synthetic data augmentation) is done with ten different random seeds, and the split between training and testing datasets is done by sampling without replacement with ten different random seeds. Spurious patterns due to correlated inputs cancel out across the different XGBoost models, whereas the true correlations are reinforced by the ensembling process. The patterns that dominate the decision-making process (and the ensuing SHAP explanations) of the ensemble are therefore more likely to be biologically relevant.

The second step of the workflow consists in quantifying the importance of each ion species with respect to a given classification task. We apply SHAP to the ensemble of ten XGBoost models obtained in the previous step. For each XGBoost model, SHAP returns one global SHAP importance score per molecular species: the global SHAP importance score is obtained by taking the mean of the magnitude of the local SHAP importance scores, or Shapley values, across all pixels making up the sample. The total global importance score of an ion species, which we previously referred to as the SHAP score, is the mean of the ten global SHAP scores obtained for that species from the ten XGBoost models making up the ensemble. The SHAP score, therefore, provides an experiment-wide measure of an ion species' predictive importance with respect to a given classification task.

We translate the task of biomarker candidate discovery into a feature selection problem: the molecular species are ranked in descending order of SHAP score to facilitate the selection of a shortlist of molecular species that may be useful biomarker candidates. In the summary bubble plots, the size of each marker corresponds to the global SHAP importance score of a given molecular species (column) for a given dataset (row). The next question is whether a biomarker candidate is positively or negatively correlated with a given target characteristic. We measure the direction and magnitude of the relationship by computing the Spearman rank-order correlation coefficient $\boldsymbol{\rho}$ between the mean-centered intensity and the Shapley values of a given molecular species. The Spearman rank-order correlation coefficient $\boldsymbol{\rho}$ also provides a way of assessing the statistical significance of the relationship: $\boldsymbol{\rho}$ ranges from -1 to 1, and we consider $\boldsymbol{\rho}$ to be significant if its magnitude exceeds 0.2. In the bar charts and summary bubble plots, the marker color corresponds to the Spearman rank-order correlation coefficient per molecular species (column) and per dataset (row).

**Table S1. Product Information for Antibodies.**

| Target | Company | Species | Clone | Lot | Cell Types/Structures | RRID | Catalog Number | Fluorophore |
|---|---|---|---|---|---|---|---|---|
| Collagen IV (α1/2) | Millipore Sigma | Mouse | 7S Domain | 3953676 | Tubular Basement Membrane, Mesangial Matrix, and Glomerular Capsule | AB_2229703 | MAB3326 | None (uses secondary) |
| Collagen IV (α5) | Cosmo Bio USA | Rat | B51 | 002 | Glomerular Basement Membrane, Glomerular Capsule, and Collecting Duct and Distal Convoluted Tubule Basement Membrane | AB_3065220 | SGE-CFT451 | FITC (did not work, used secondary) |
| Tensin | Sigma-Aldrich | Rabbit | TNS1 | A115685 | Mesangial Cells and Vascular Smooth Muscle Cells | AB_10796111 | HPA036089 | None (uses secondary) |
| Podocalyxin | Abcam | Rabbit | PODXL | 1000363-1 | Podocyte Cytoplasm and Plasma Membrane | AB_3065225 | AB208254 | AF 488 |
| CD31 | Abcam | Mouse | JC/7OA | 1038052-1 | Endothelial Cells | AB_2890260 | AB215912 | AF 647 |
| Fibronectin | Abcam | Rabbit | Fibronectin | 1054792-1 | Mesangial Matrix and Muscularized Vessel Walls | AB_3086599 | AB309604 | AF 594 |
| Synaptopodin | Abcam | Rabbit | Synaptopodin | GR3400313-3 | Podocyte Cytoplasm and Plasma Membrane | AB_3086600 | AB282117 | Cy7 (conjugated with Lightning Link) |
| Nestin | Novus | Mouse | 10C2 | D127147 | Podocyte Cytoplasm | AB_922042 | NB300-266AF488 | AF 488 |
| αSMA | Abcam | Mouse | 1A4 | GR3395316-5 | Vascular Smooth Muscle Cells | AB_2924381 | AB202368 | AF 594 |
| AQP1 | Abcam | Rabbit | Aquaporin 1 | 1028514-1 | Proximal Tubules | AB_3086601 | AB225225 | AF 647 |
| Secondary anti-rat | Jackson Immuno Research | Donkey | X | 161323 | X | AB_2340667 | 712-165-153 | Cy3 |
| Secondary anti-mouse | Jackson Immuno Research | Donkey | X | 160571 | X | AB_2340819 | 715-175-150 | Cy5 |
| Secondary anti-rabbit | Jackson Immuno Research | Donkey | X | 162188 | X | AB_2313584 | 711-545-152 | AF488 |

**Table S2. Instrument Parameters for Positive Ion Mode qTOF MALDI IMS Experiments.**

| **Positive Ion Mode** | |
|---|---|
| Transfer | |
| MALDI Plate Offset | 70.0 V |
| Deflection 1 Delta | 70.0 V |
| Funnel 1 RF | 400.0 Vpp |
| isCID Energy | 5.0 eV Vpp |
| Funnel 2 RF | 350.0 Vpp |
| Multipole RF | 400.0 Vpp |
| Collision Cell | |
| Collision Energy | 10.0 eV |
| Collision RF | 2000.0 Vpp |
| Quadrupole | |
| Ion Energy | 10.0 eV |
| Low Mass | *m/z* 450.00 |
| Focus Pre TOF | |
| Transfer Time | 100 µs |
| Pre Pulse Storage | 10.0 µs |

**Table S3. Instrument Parameters for Negative Ion Mode LC-MS/MS Experiments.**

| **Negative Ion Mode** | | |
|---|---|---|
| ESI | | |
| Capillary | 3600 V | |
| Nebulizer | 2.0 Bar | |
| Dry Gas | 8.0 l/min | |
| Dry Temp | 220 °C | |
| MS/MS Stepping | | |
| $1/K_0$ | Scan 1 | Scan 2 |
| 0.40 | 40 eV | 40 eV |
| 2.32 | 50 eV | 65 eV |
| Collision RF | 450 Vpp | 1800 Vpp |
| Transfer Time | 25 µs | 120 µs |
| Pre Pulse Storage | 5 µs | 10 µs |

**Table S4. Instrument Parameters for Positive Ion Mode LC-MS/MS Experiments.**

| **Positive Ion Mode** | | |
|---|---|---|
| ESI | | |
| Capillary | 4500 V | |
| Nebulizer | 2.0 Bar | |
| Dry Gas | 8.0 l/min | |
| Dry Temp | 220 °C | |
| MS/MS Stepping | | |
| $1/K_0$ | Scan 1 | Scan 2 |
| 0.50 | 40 eV | 40 eV |
| 1.90 | 40 eV | 40 eV |
| Collision RF | 500 Vpp | 1500 Vpp |
| Transfer Time | 25 µs | 85 µs |
| Pre Pulse Storage | 5 µs | 8 µs |

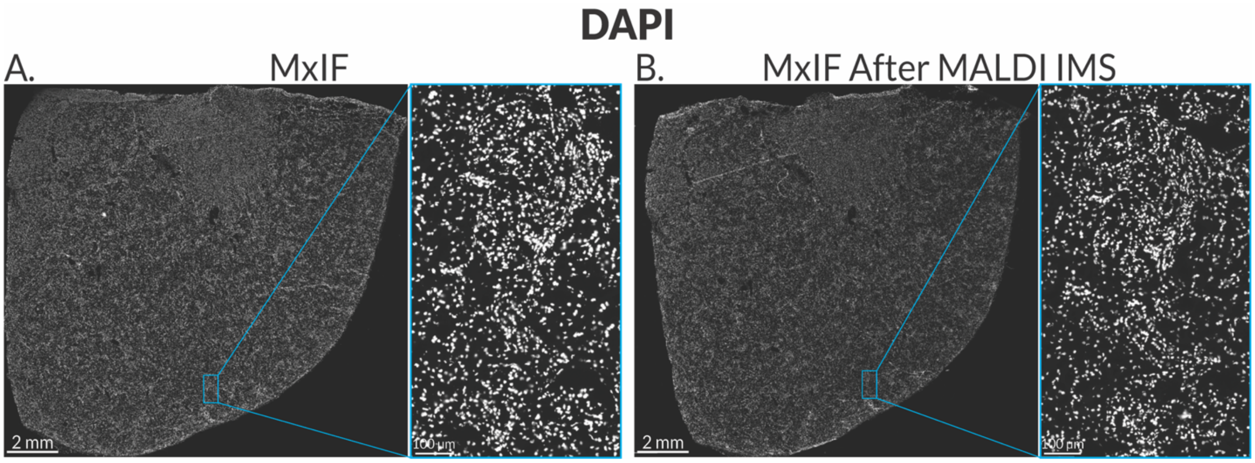


**Figure S1.** Grayscale comparison of the DAPI stain from a tissue section that was only stained with MxIF (**A**) and a tissue section that was imaged with MALDI IMS first and then stained with MxIF (**B**).

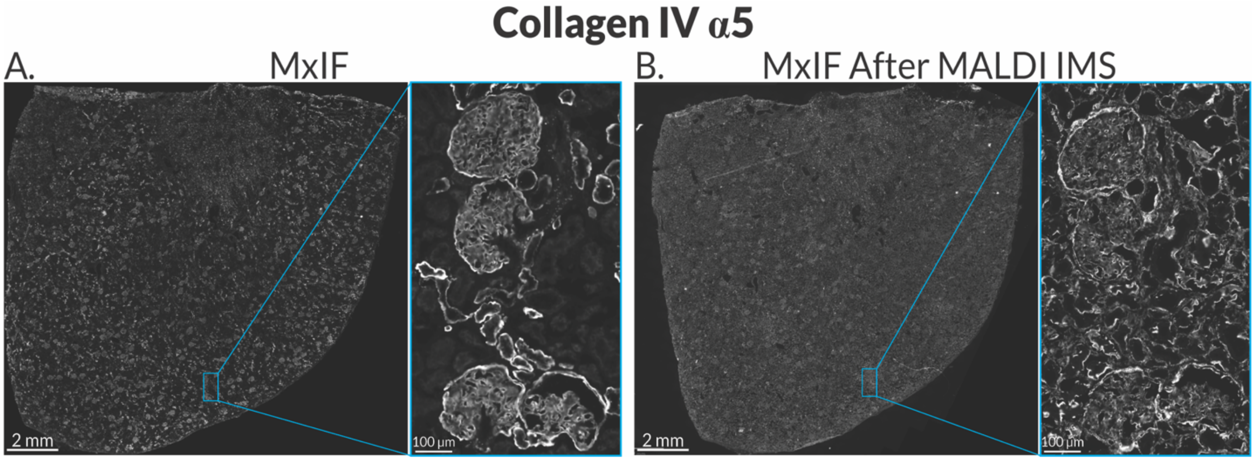


**Figure S2.** Grayscale comparison of the collagen IV α5 stain from a tissue section that was only stained with MxIF (**A**) and a tissue section that was imaged with MALDI IMS first and then stained with MxIF (**B**).

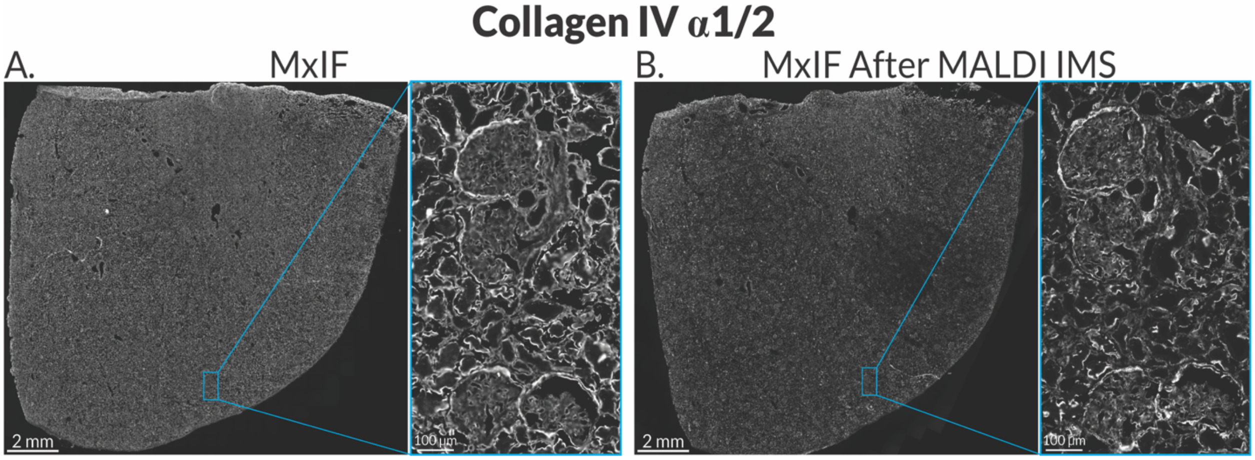


**Figure S3.** Grayscale comparison of the collagen IV α1/2 stain from a tissue section that was only stained with MxIF (**A**) and a tissue section that was imaged with MALDI IMS first and then stained with MxIF (**B**).

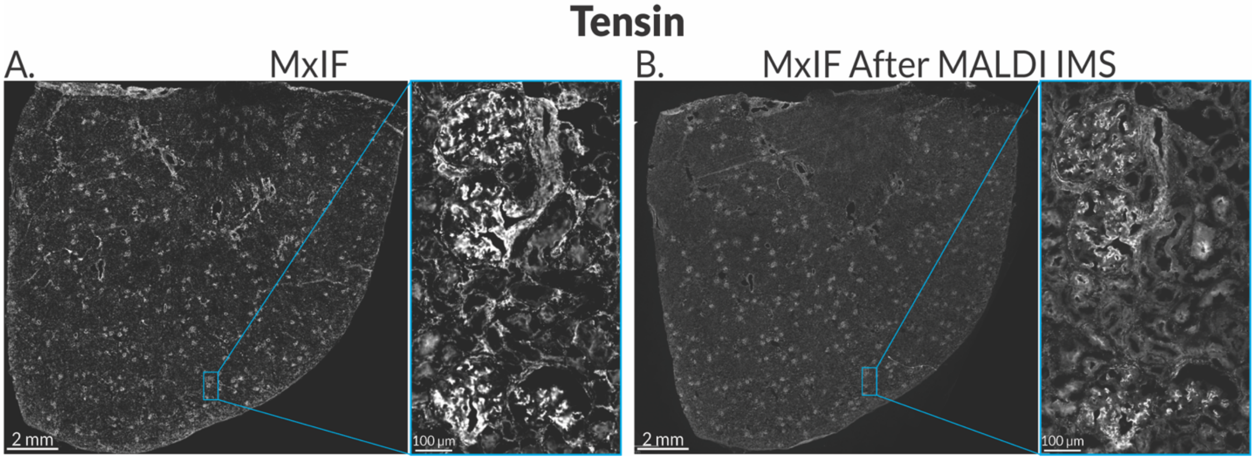


**Figure S4.** Grayscale comparison of the tensin stain from a tissue section that was only stained with MxIF (**A**) and a tissue section that was imaged with MALDI IMS first and then stained with MxIF (**B**).

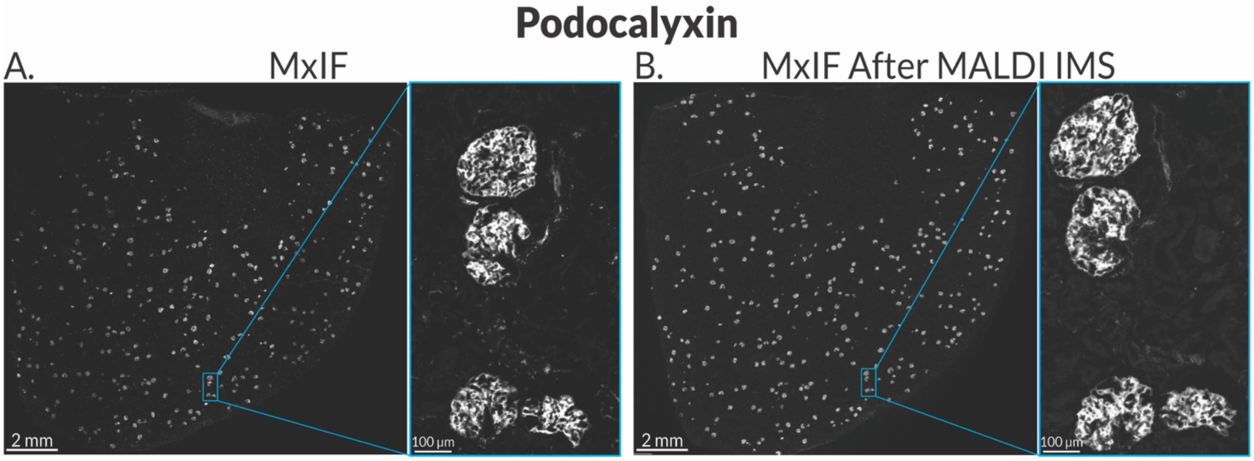


**Figure S5.** Grayscale comparison of the podocalyxin stain from a tissue section that was only stained with MxIF (**A**) and a tissue section that was imaged with MALDI IMS first and then stained with MxIF (**B**).

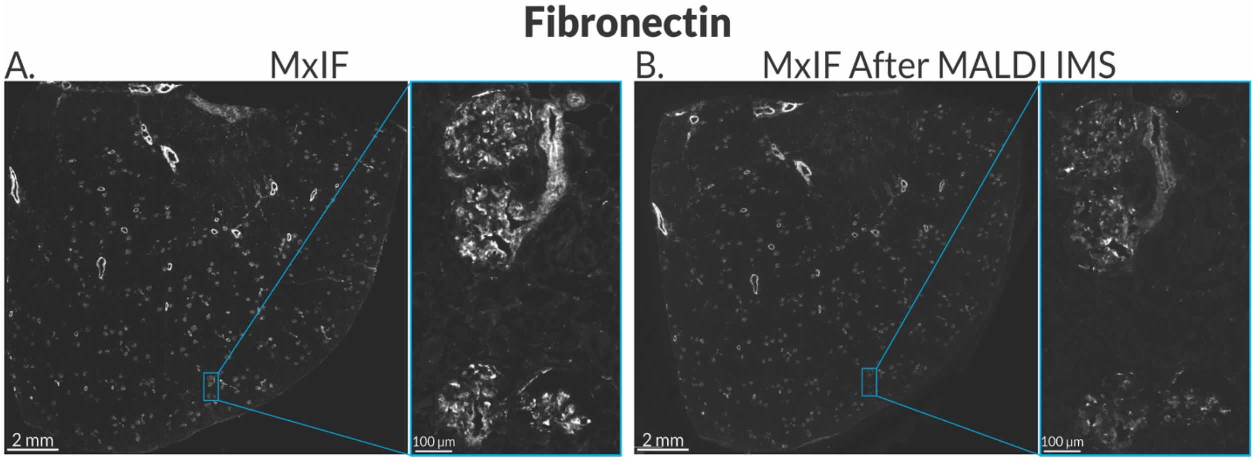


**Figure S6.** Grayscale comparison of the fibronectin stain from a tissue section that was only stained with MxIF (**A**) and a tissue section that was imaged with MALDI IMS first and then stained with MxIF (**B**).

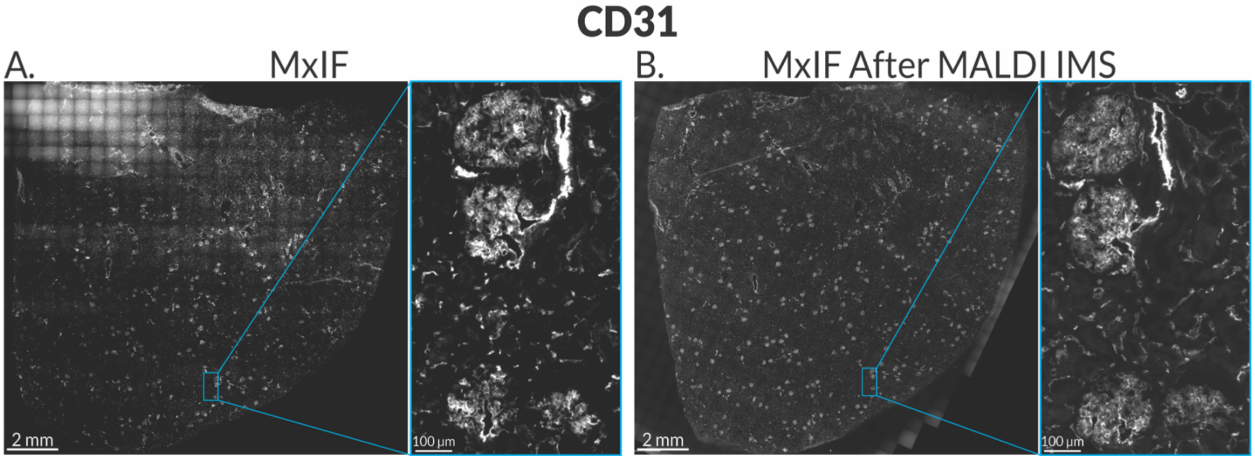


**Figure S7.** Grayscale comparison of the CD31 stain from a tissue section that was only stained with MxIF (**A**) and a tissue section that was imaged with MALDI IMS first and then stained with MxIF (**B**).

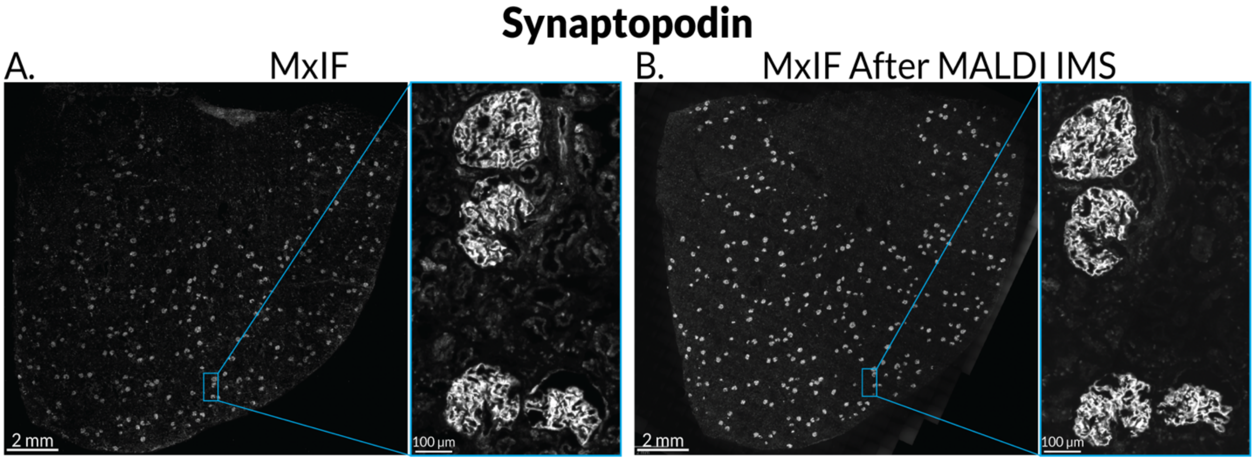


**Figure S8.** Grayscale comparison of the synaptopodin stain from a tissue section that was only stained with MxIF (**A**) and a tissue section that was imaged with MALDI IMS first and then stained with MxIF (**B**).

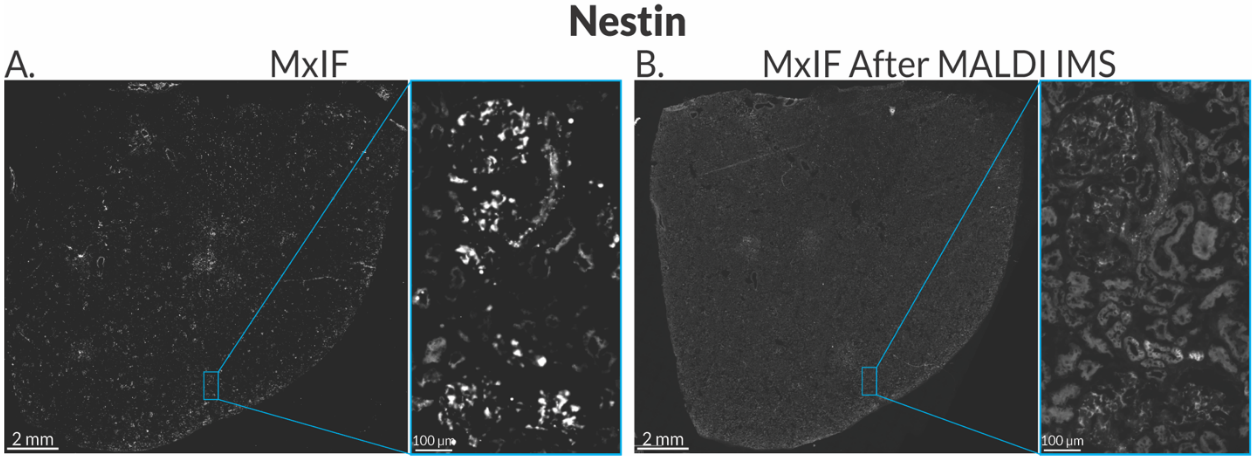


**Figure S9.** Grayscale comparison of the nestin stain from a tissue section that was only stained with MxIF (**A**) and a tissue section that was imaged with MALDI IMS first and then stained with MxIF (**B**).

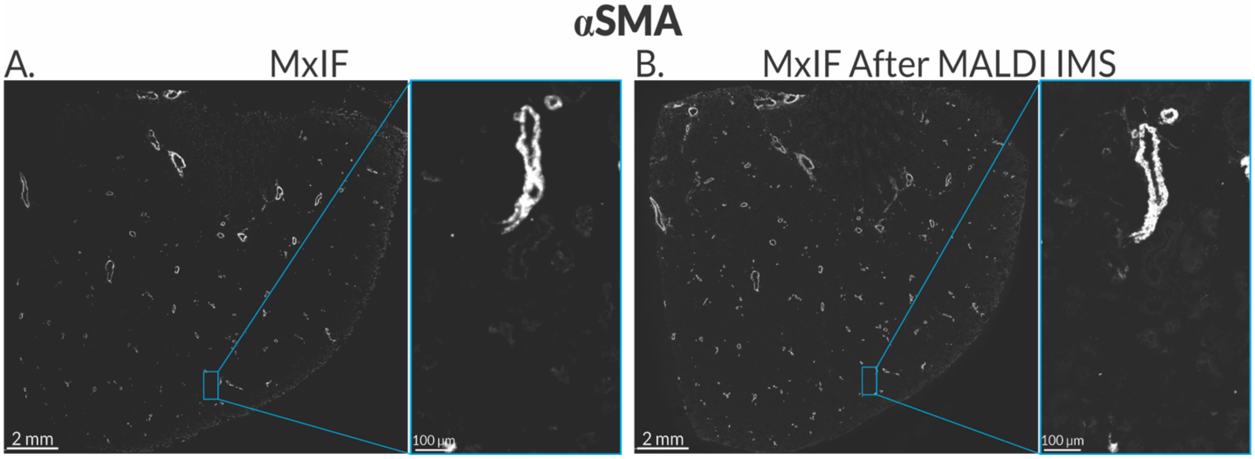


**Figure S10.** Grayscale comparison of the αSMA stain from a tissue section that was only stained with MxIF (**A**) and a tissue section that was imaged with MALDI IMS first and then stained with MxIF (**B**).

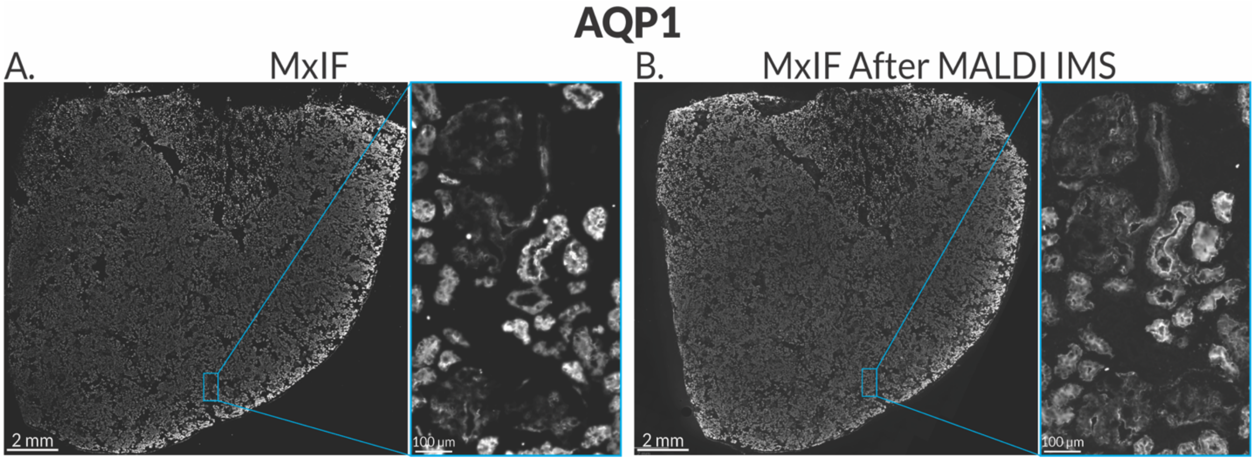


**Figure S11.** Grayscale comparison of the AQP1 stain from a tissue section that was only stained with MxIF (**A)** and a tissue section that was imaged with MALDI IMS first and then stained with MxIF (**B).**

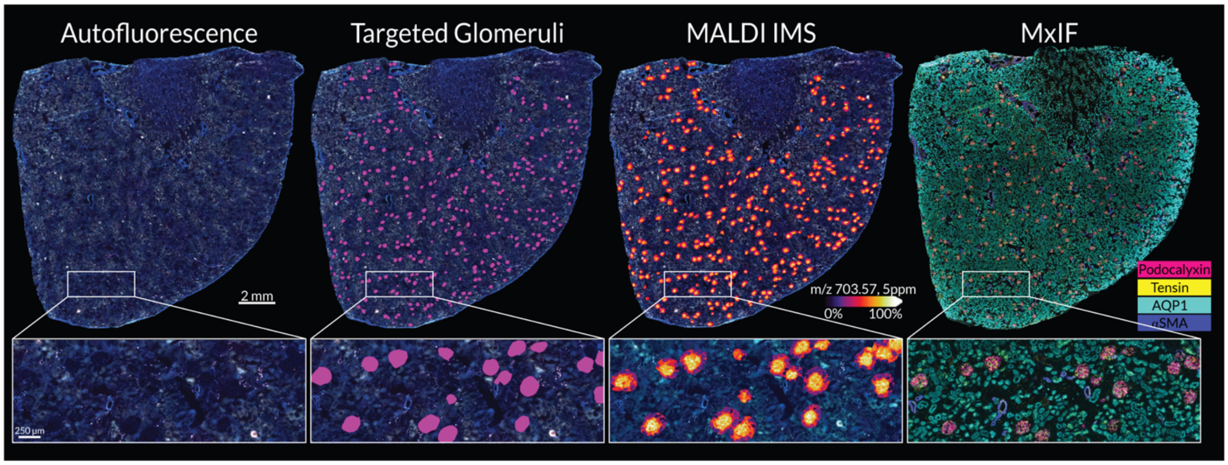


**Figure S12.** Overall imaging workflow. The whole-slide autofluorescence image of the tissue section (**A**) is used to segment glomeruli across the section (**B**). A targeted MALDI IMS experiment is performed on only glomerular regions (**C**). After the MALDI IMS experiment, the matrix can be removed and MxIF can be stained on the same tissue section.

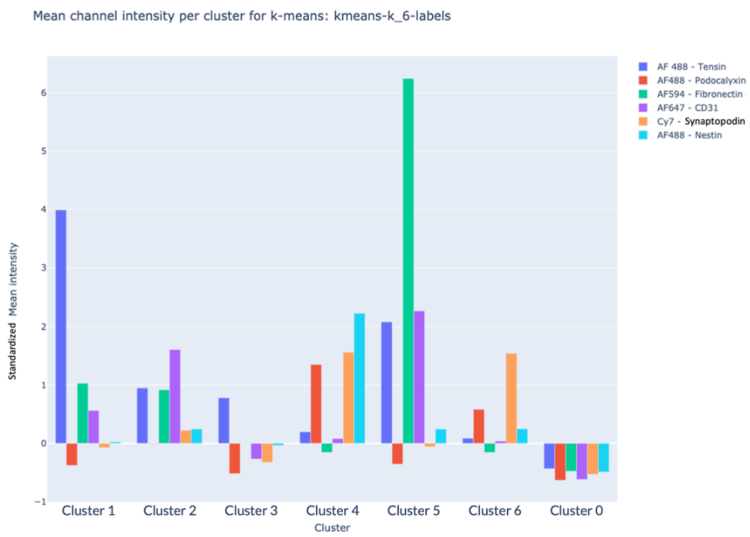


**Figure S13.** Plot of standardized mean fluorescent intensities of antibodies for each *k*-means cluster. Note, a non-cell specific cluster (cluster 3) includes tissue regions with low-level signal from tensin and minimal MxIF signal detected in other channels. These regions include pixels with seemingly high background signal and low cell type specificity. A zero/low MxIF intensity cluster (cluster 0) captures regions with minimal MxIF signal for this antibody marker panel. These low MxIF-signal regions may include tissue features not stained using the included antibodies.

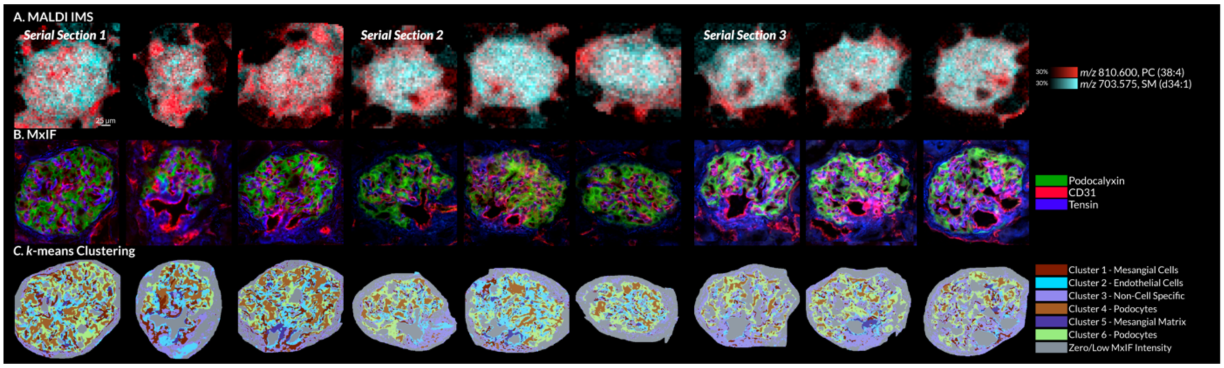

**Figure S14. Multimodal molecular imaging data and segmentation maps of nine example glomeruli from three serial tissue sections.** MALDI-based ion images for PC(38:4) (*m/z* 810.600) and SM(d34:1) (*m/z* 703.575) (**A**). Overlaid MxIF images of podocalyxin, CD31, and tensin (**B**). All *k*-means clustering-based segments of the MxIF data and the glomerular cell types that tend to dominate each segment (**C**).

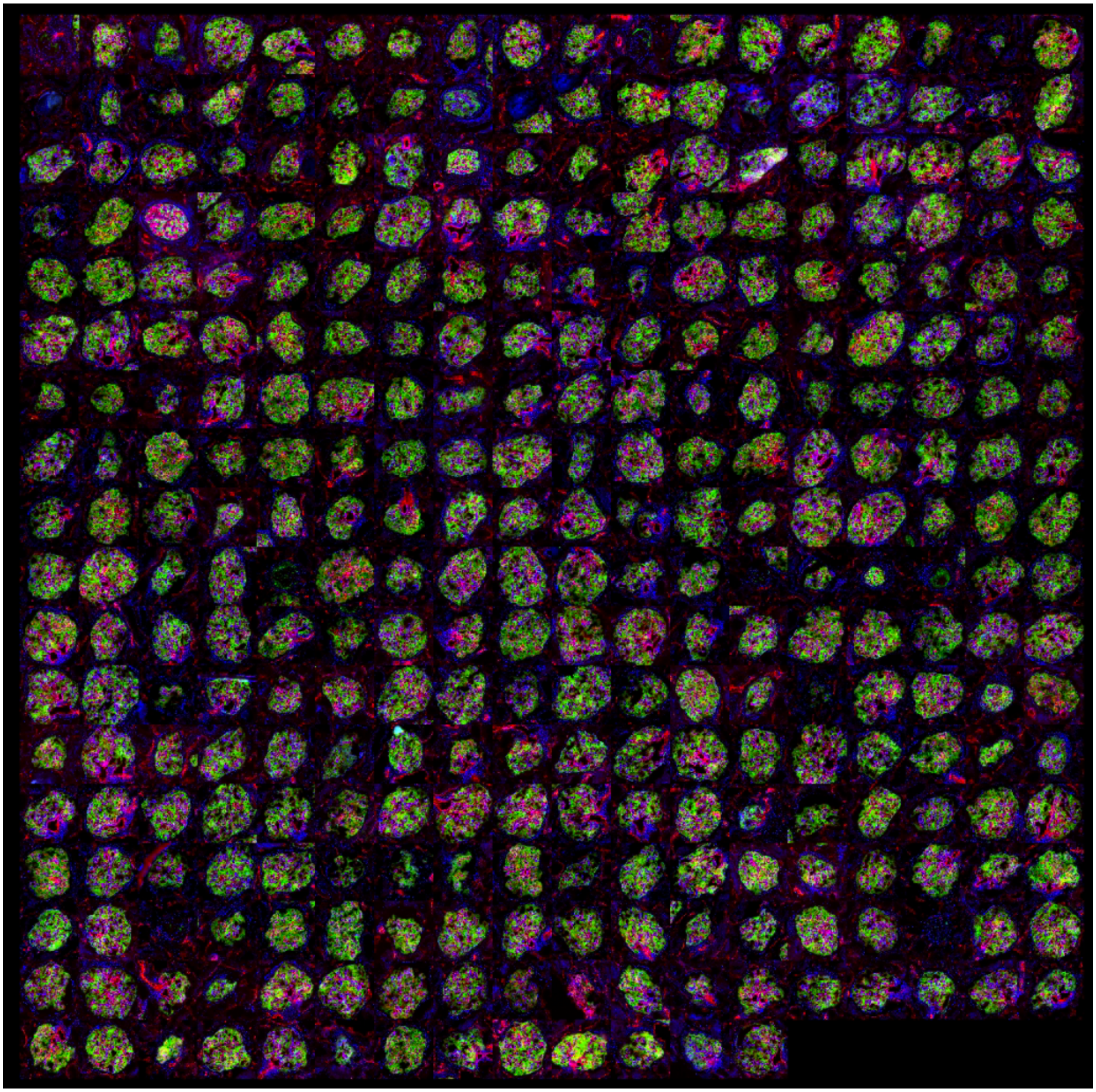

**Figure S15.** MxIF mosaic of all glomeruli from the first of the triplicate serial sections. In this mosaic, podocalyxin is represented in green, CD31 is represented in red, and tensin is represented in blue.

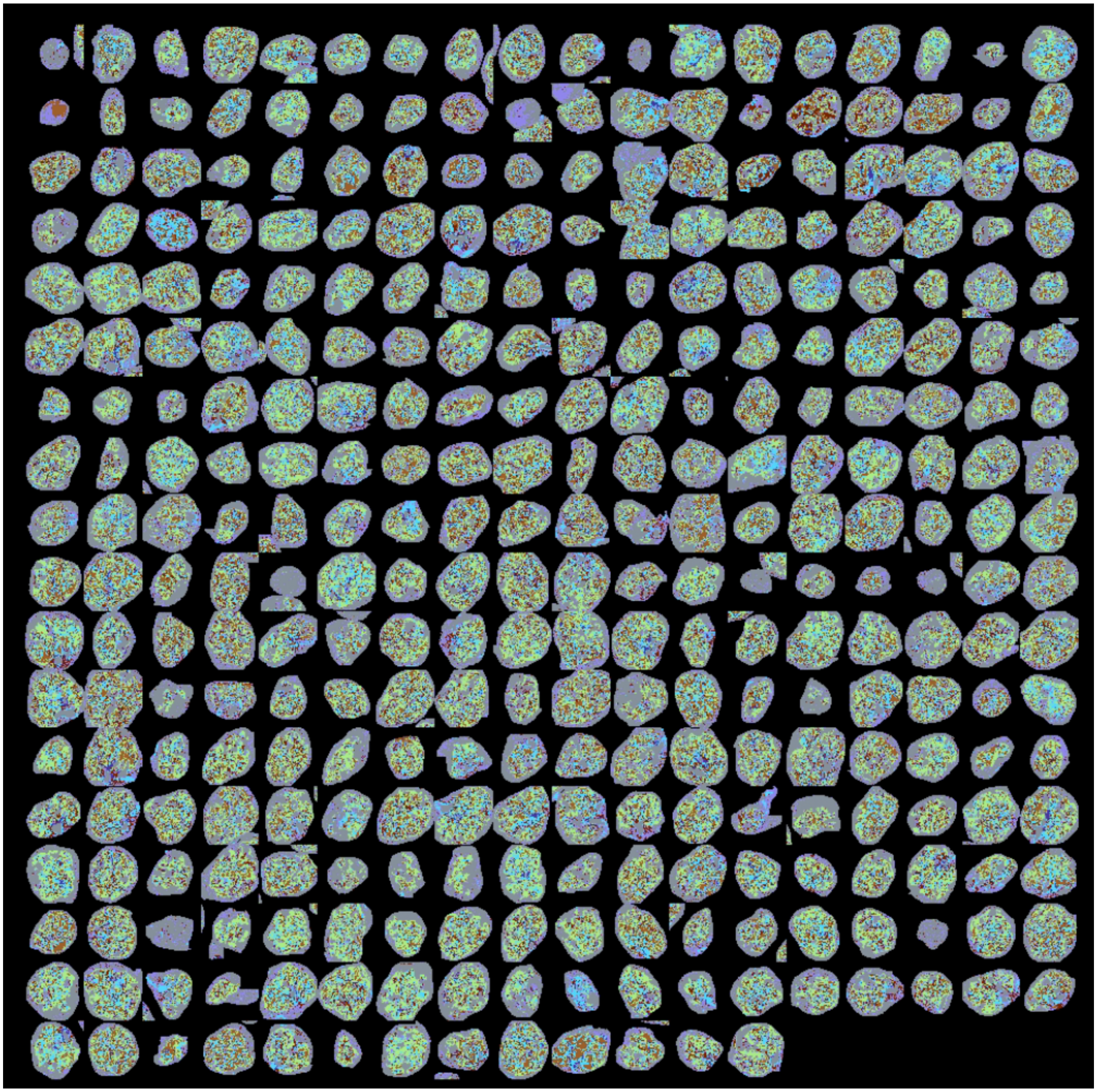

**Figure S16.** *k*-means glomerular clusters mosaic of all glomeruli from the first of the triplicate serial sections. In this mosaic, the clusters are represented by the same colors as reported in Figure 2.

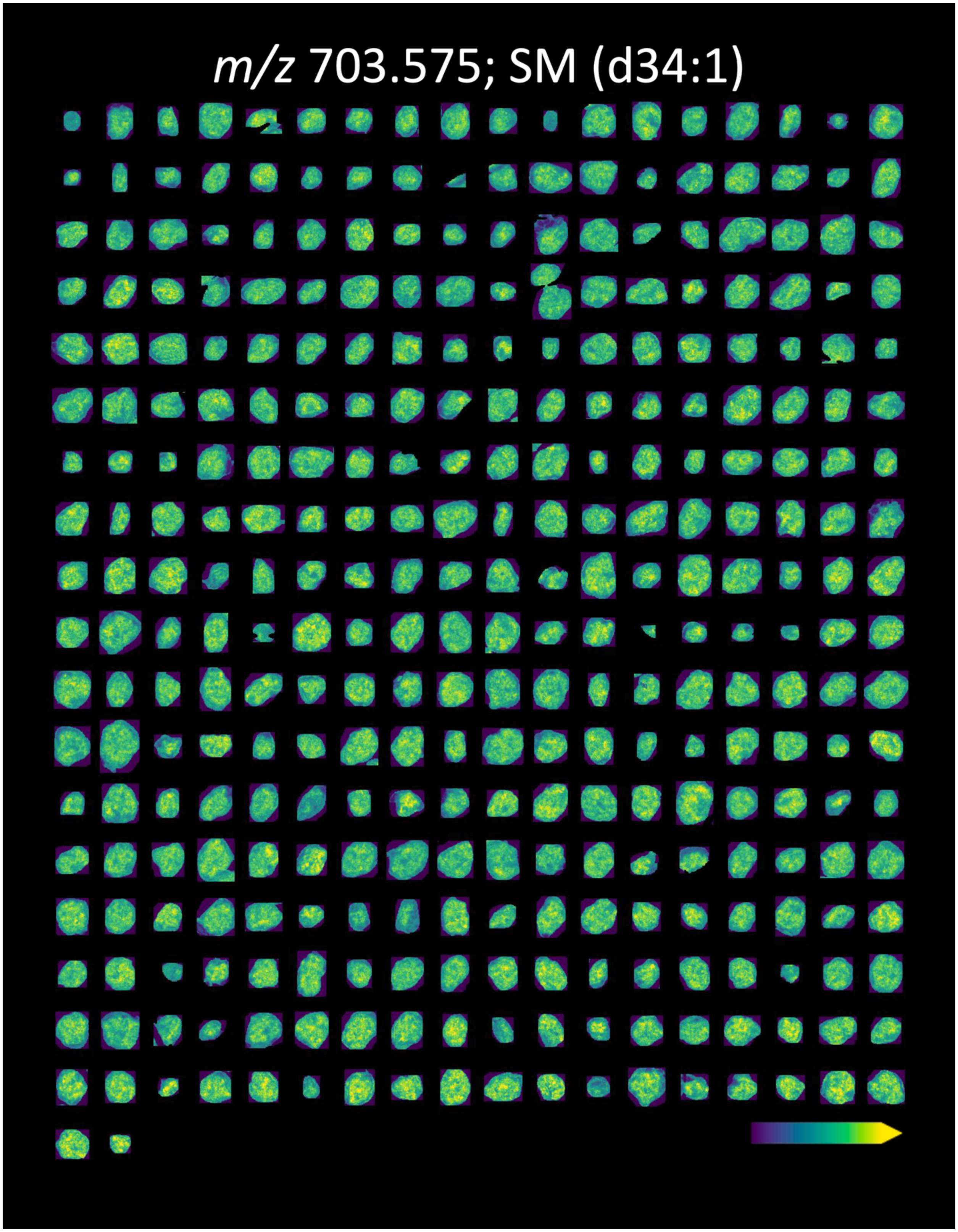


**Figure S17.** Mosaic ion image of all glomeruli from the first of the triplicate serial sections. SM (d34:1) has a mass error of 0.28 ppm.

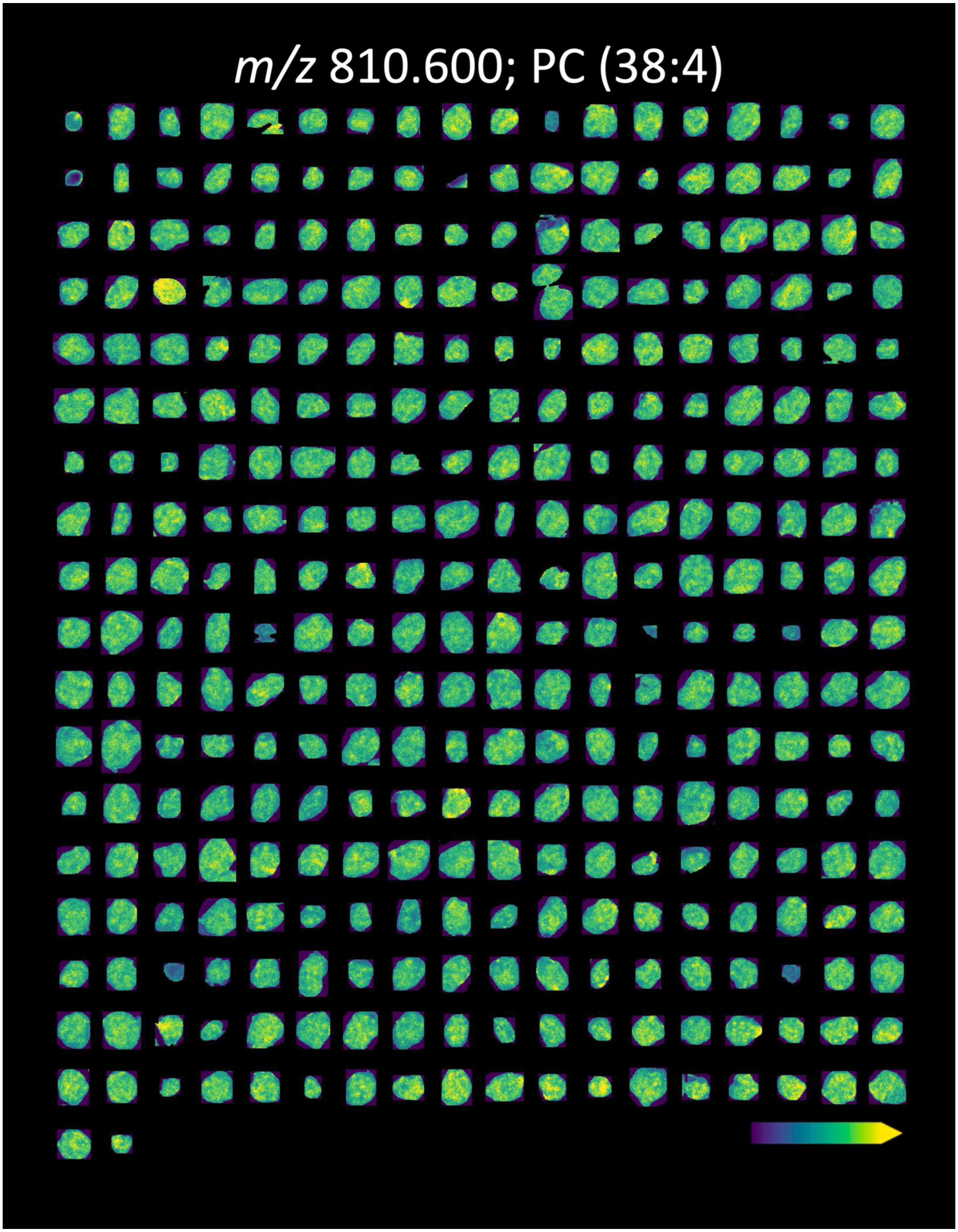


**Figure S18.** Mosaic ion image of all glomeruli from the first of the triplicate serial sections. PC (38:4) has a mass error of -0.86 ppm.

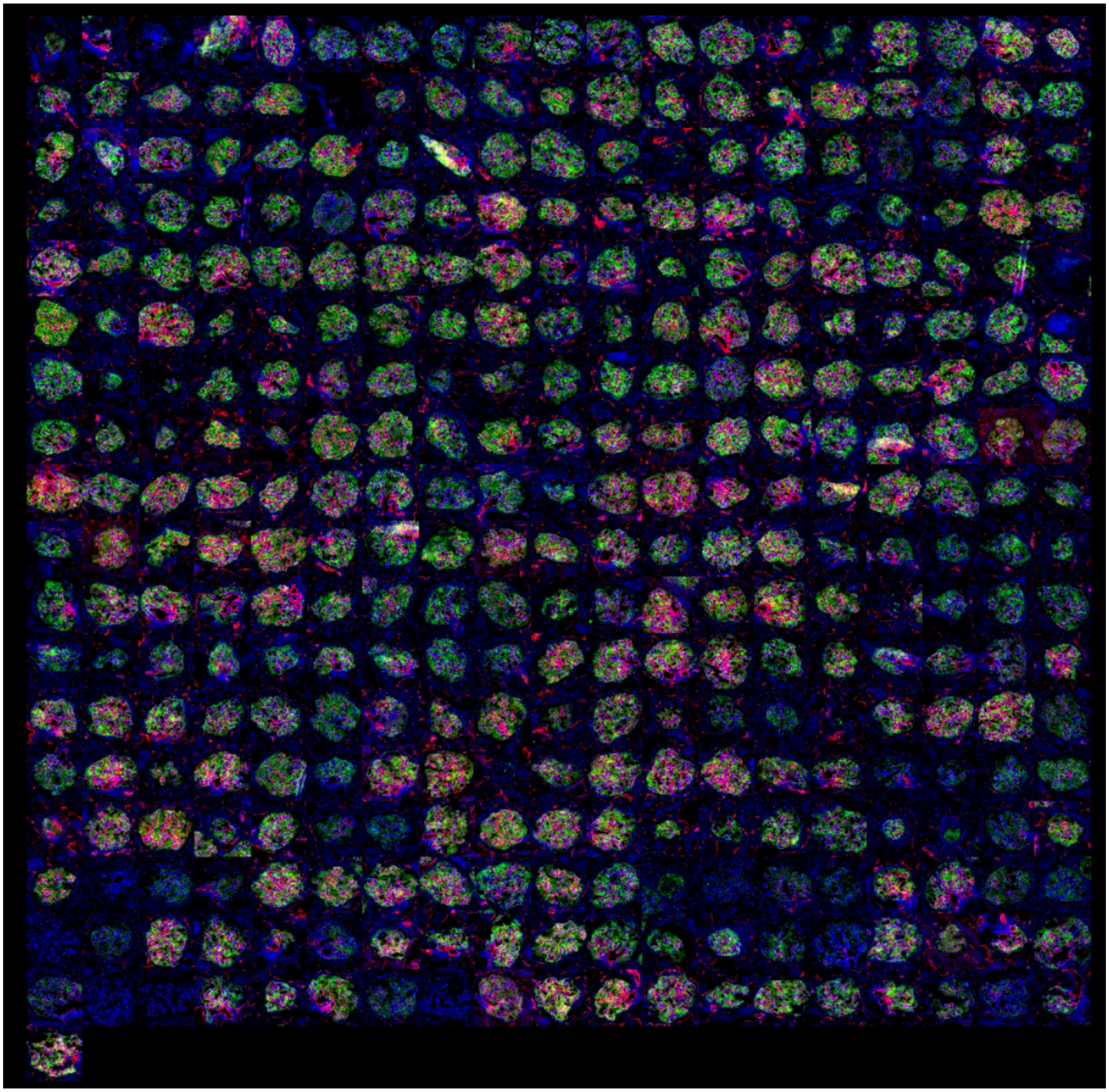

**Figure S19.** MxIF mosaic of all glomeruli from the second of the triplicate serial sections. In this mosaic, podocalyxin is represented in green, CD31 is represented in red, and tensin is represented in blue.

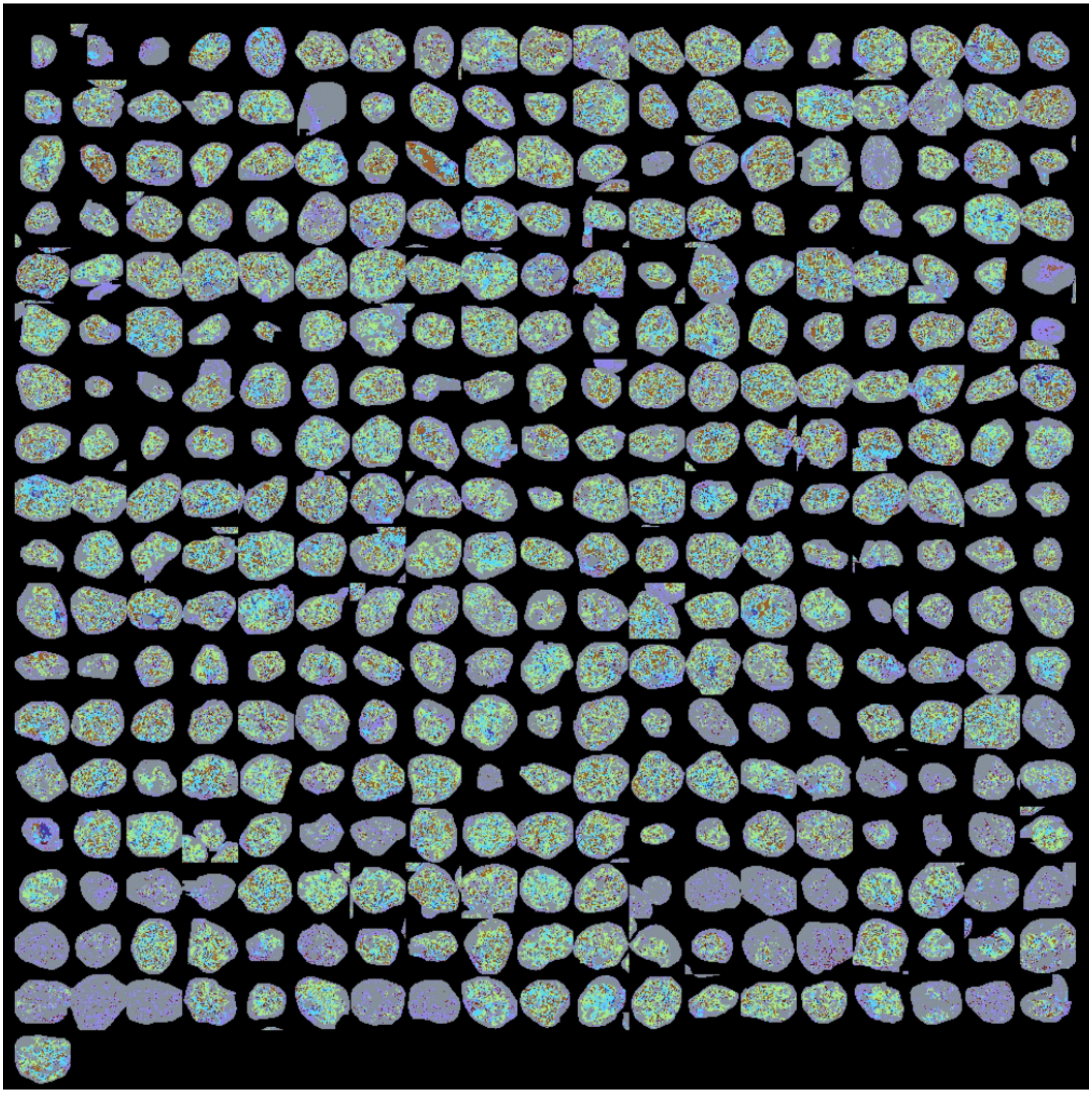

**Figure S20.** *k*-means glomerular clusters mosaic of all glomeruli from the second of the triplicate serial sections. In this mosaic, the clusters are represented by the same colors as reported in Figure 2.

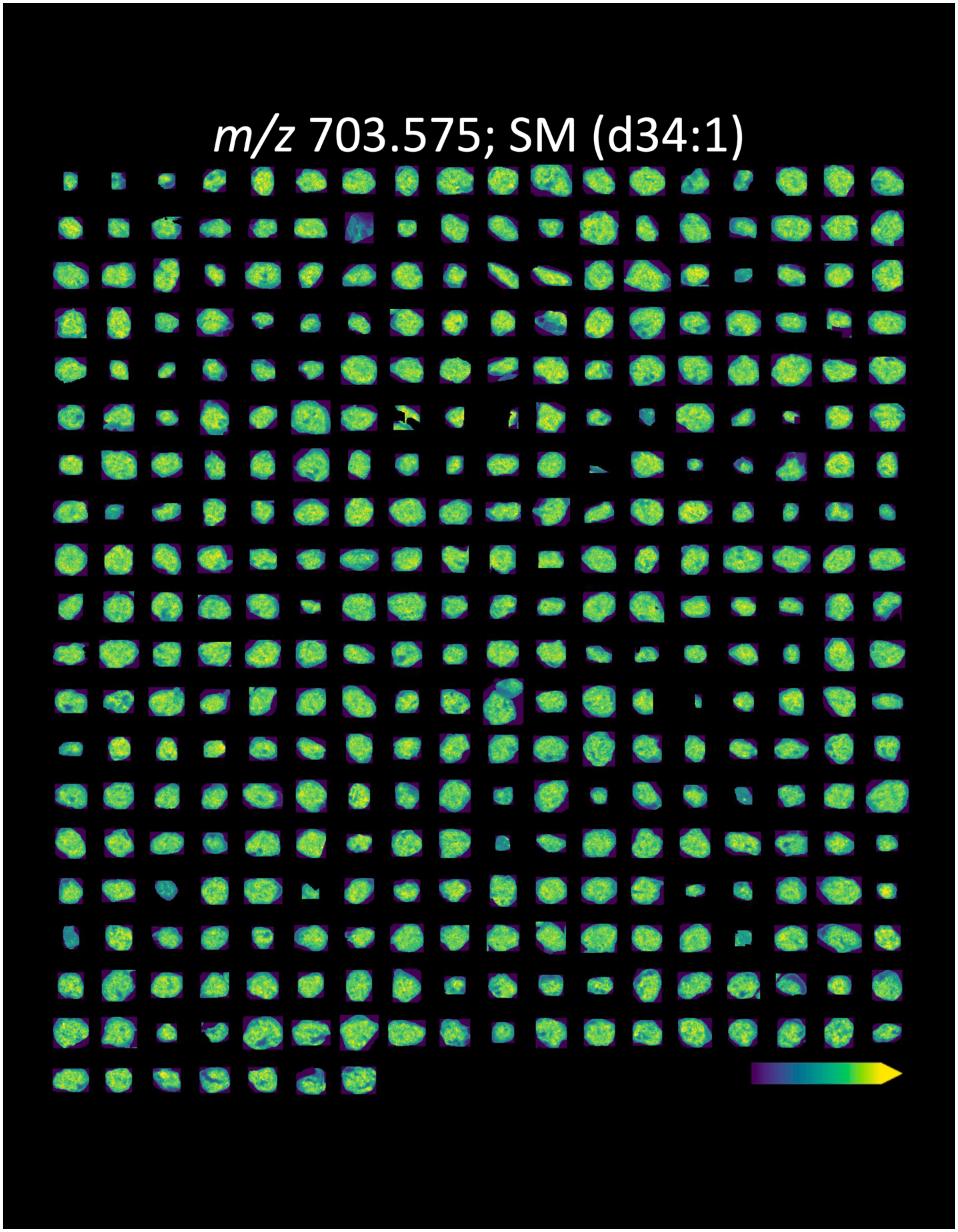


**Figure S21.** Mosaic ion image of all glomeruli from the second of the triplicate serial sections. SM (d34:1) has a mass error of 0.28 ppm.

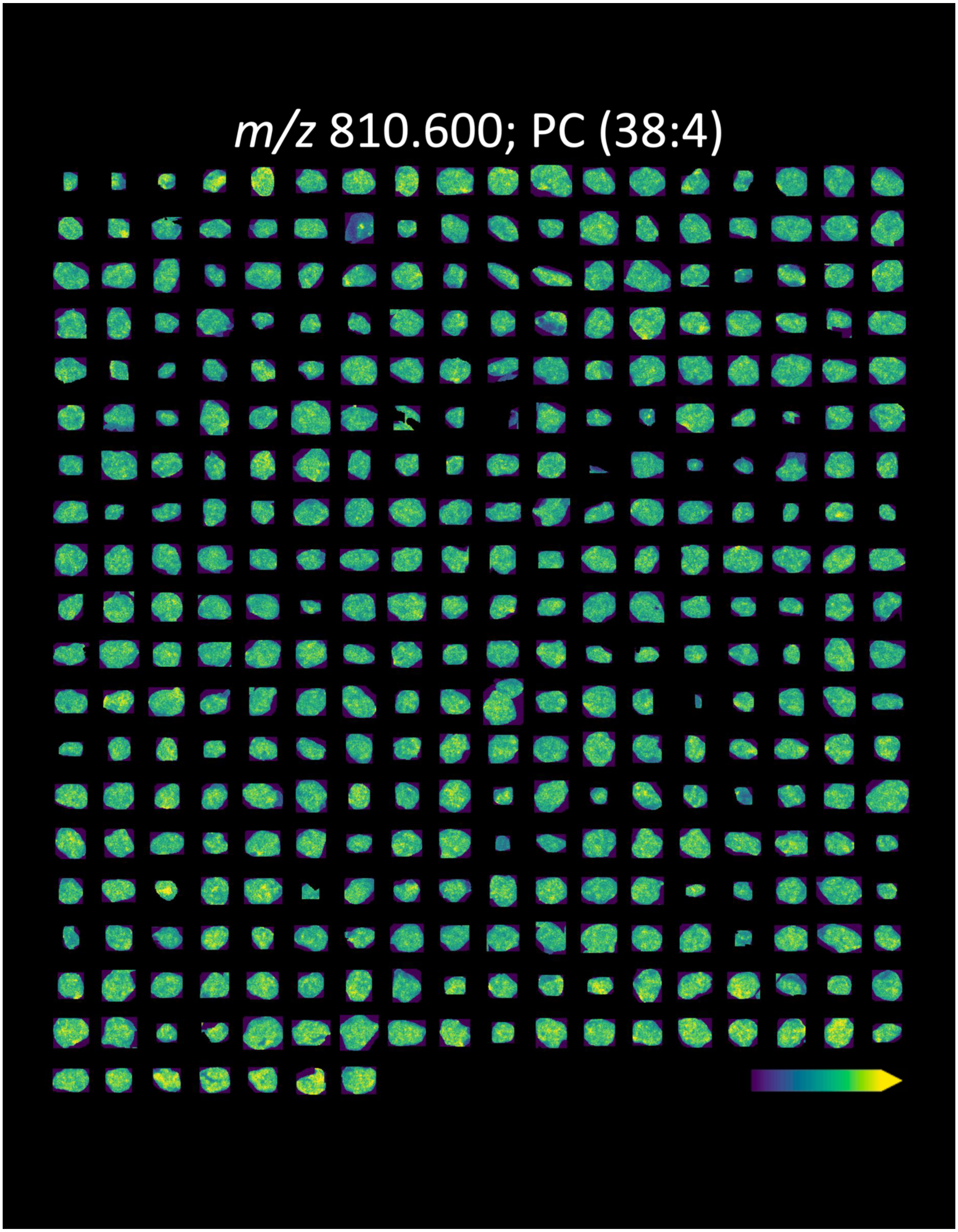


**Figure S22.** Mosaic ion image of all glomeruli from the second of the triplicate serial sections. PC (38:4) has a mass error of -0.86 ppm.

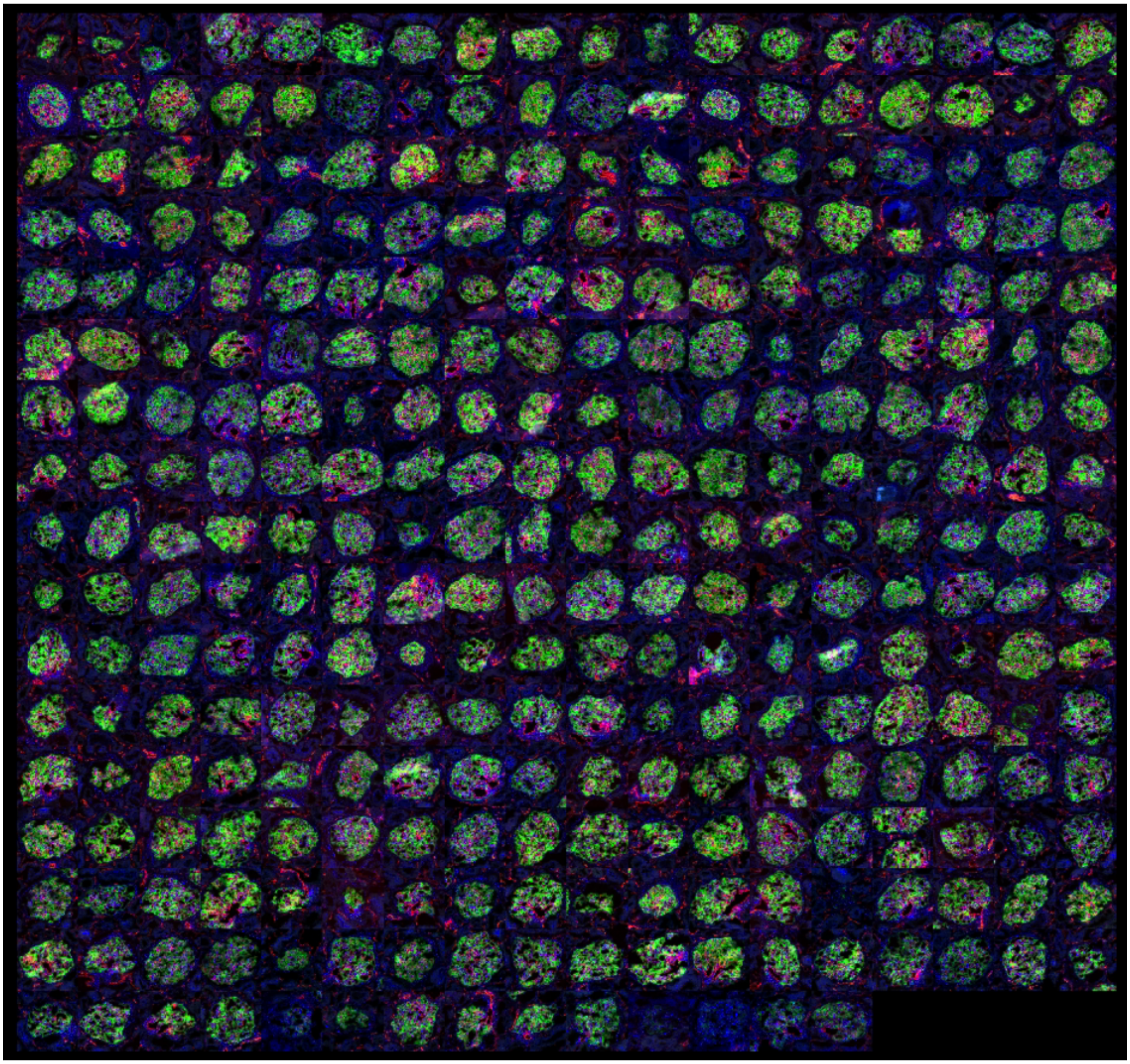

**Figure S23.** MxIF mosaic of all glomeruli from the third of the triplicate serial sections. In this mosaic, podocalyxin is represented in green, CD31 is represented in red, and tensin is represented in blue.

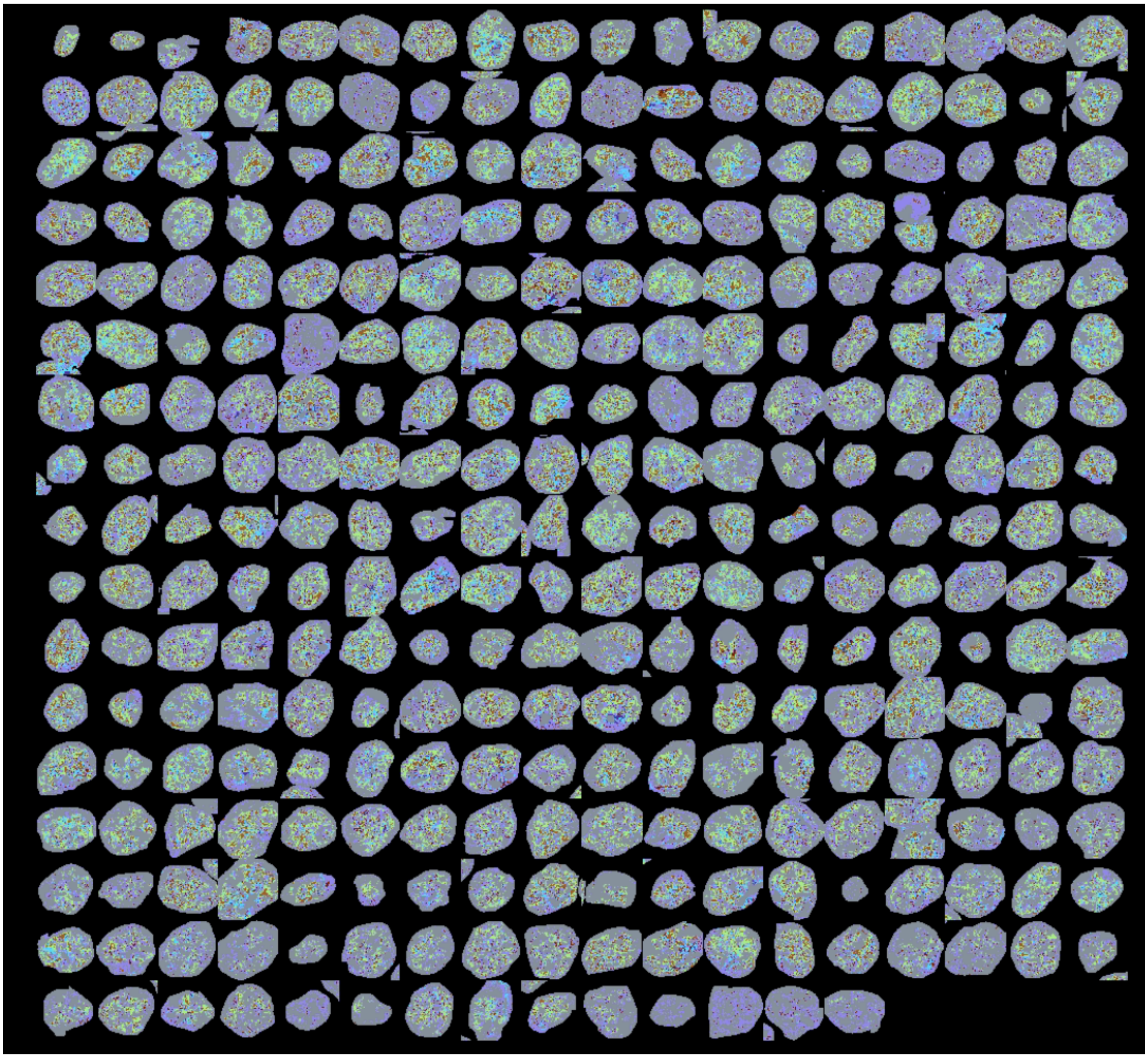

**Figure S24.** *k*-means glomerular clusters mosaic of all glomeruli from the third of the triplicate serial sections. In this mosaic, the clusters are represented by the same colors as reported in Figure 2.

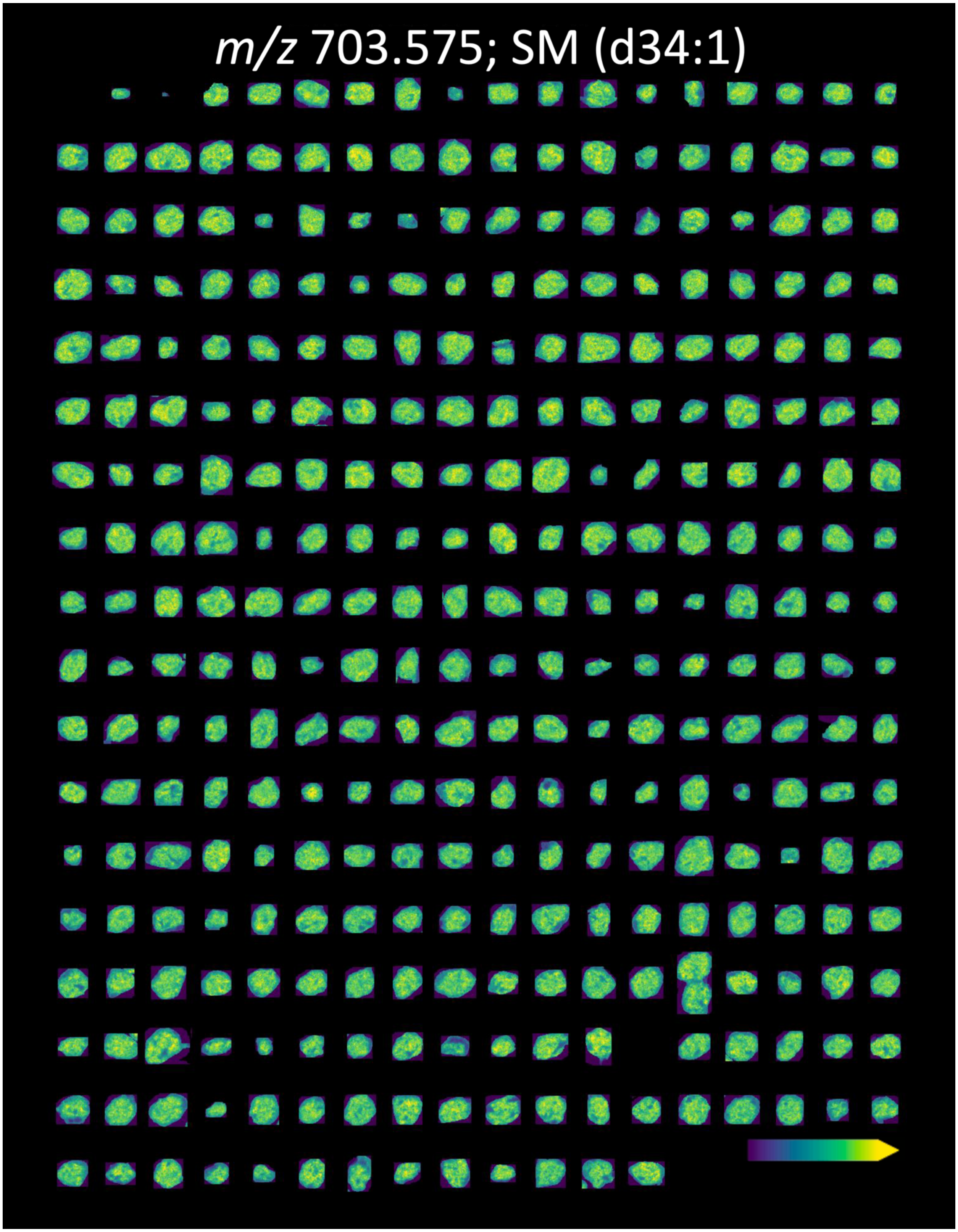

**Figure S25.** Mosaic ion image of all glomeruli from the third of the triplicate serial sections. SM (d34:1) has a mass error of 0.28 ppm.

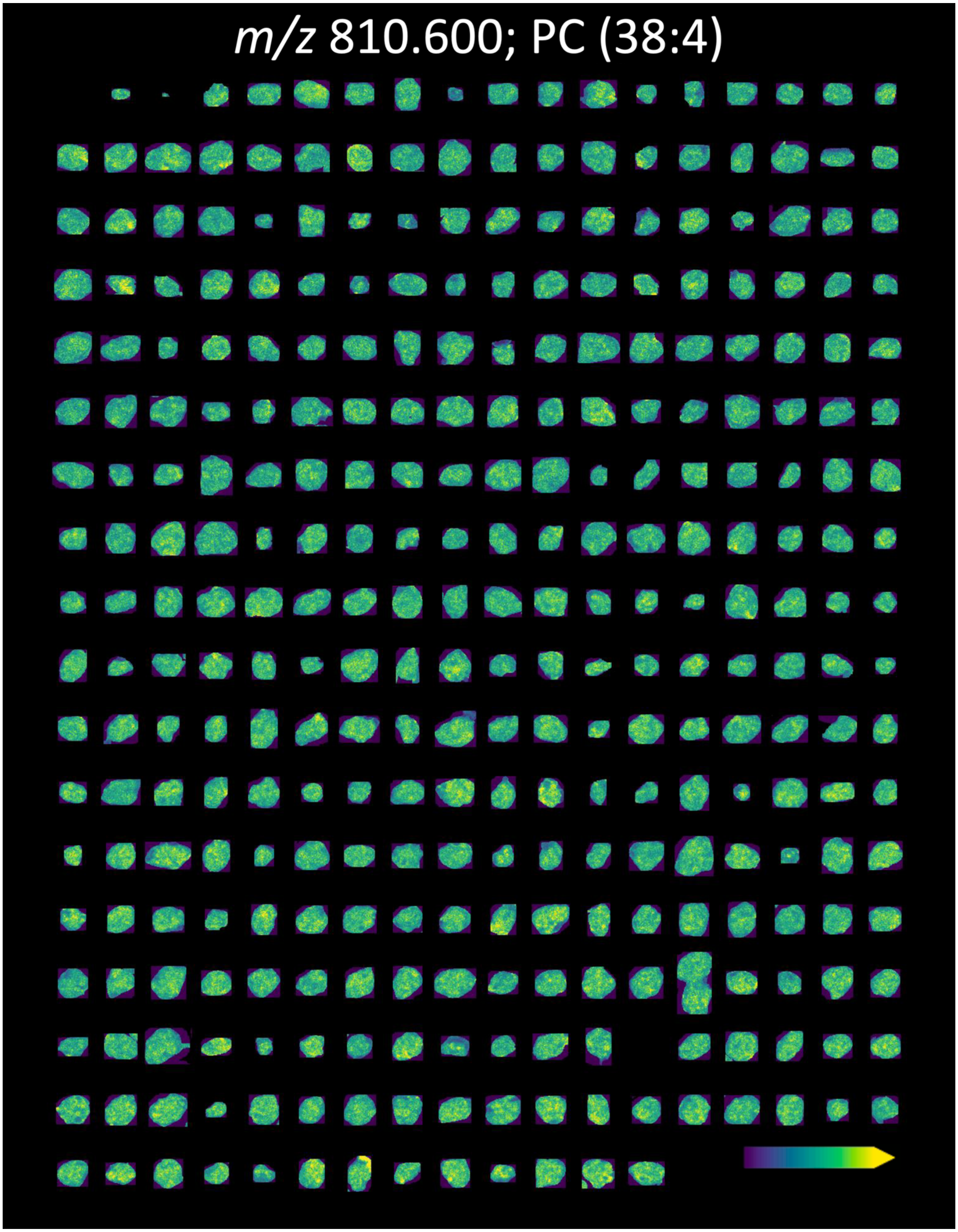


**Figure S26.** Mosaic ion image of all glomeruli from the third of the triplicate serial sections. PC (38:4) has a mass error of -0.86 ppm.

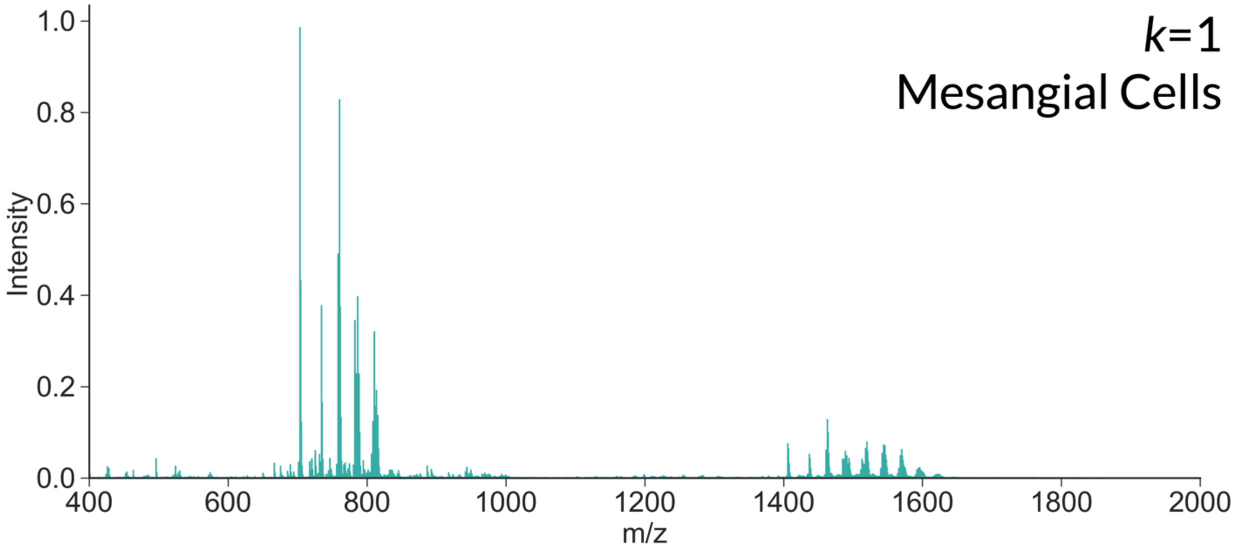


**Figure S27.** Average mass spectrum of cluster 1 (mesangial cell-related segment).

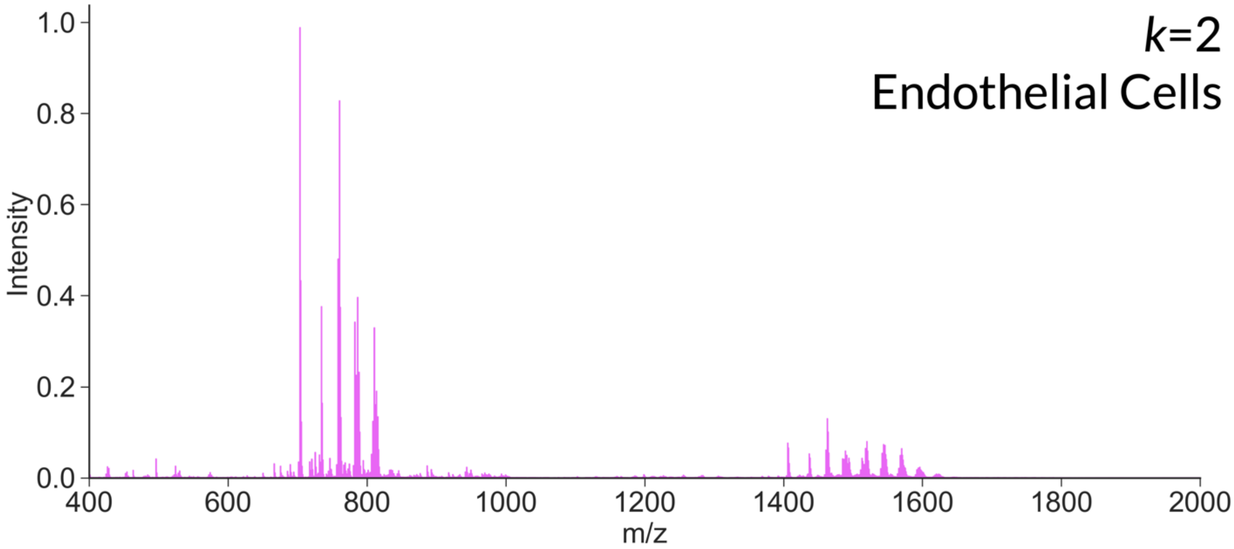


**Figure S28.** Average mass spectrum of cluster 2 (endothelial cell-related segment).

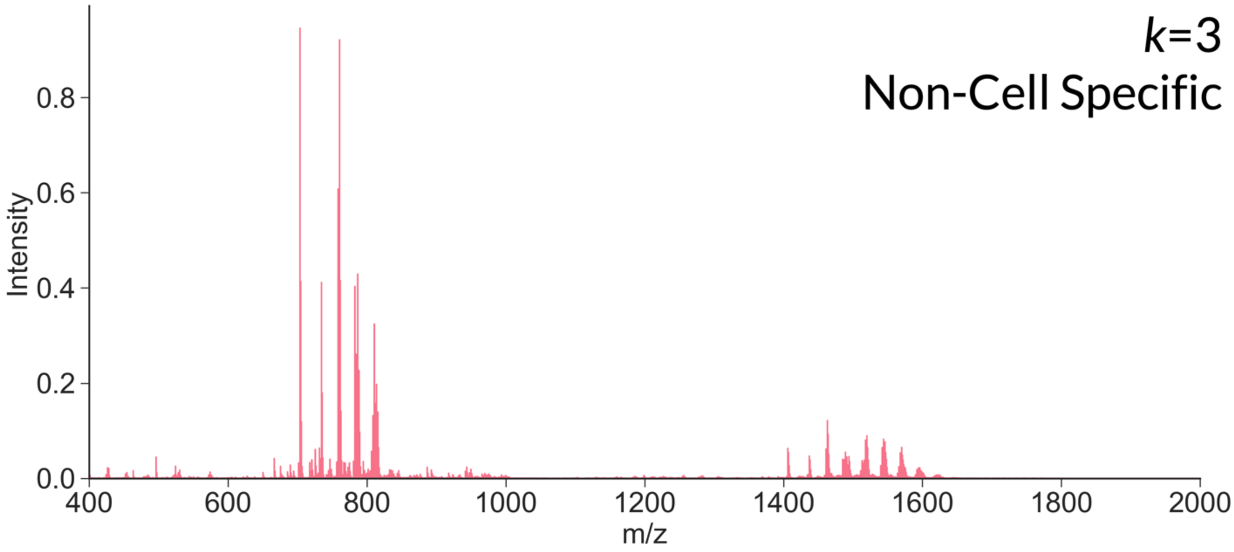


**Figure S29.** Average mass spectrum of cluster 3. This cluster is defined as regions with low-level signal from tensin and minimal MxIF signal detected in other channels. These regions include pixels with seemingly high background signal and low cell type specificity.

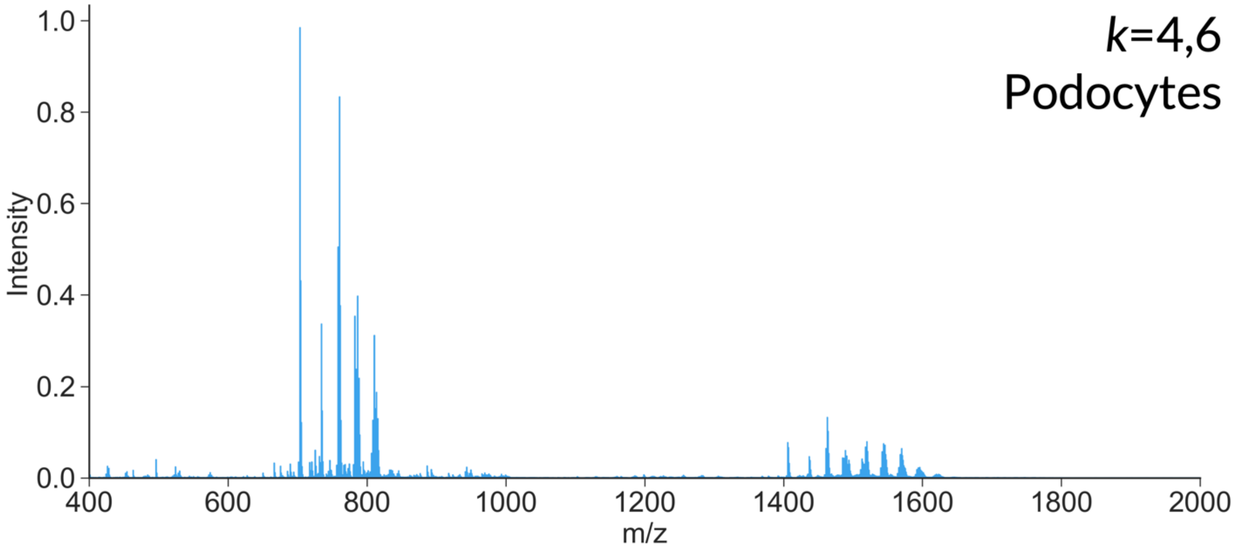


**Figure S30.** Average mass spectrum of clusters 4 and 6 combined (podocyte-related segment).

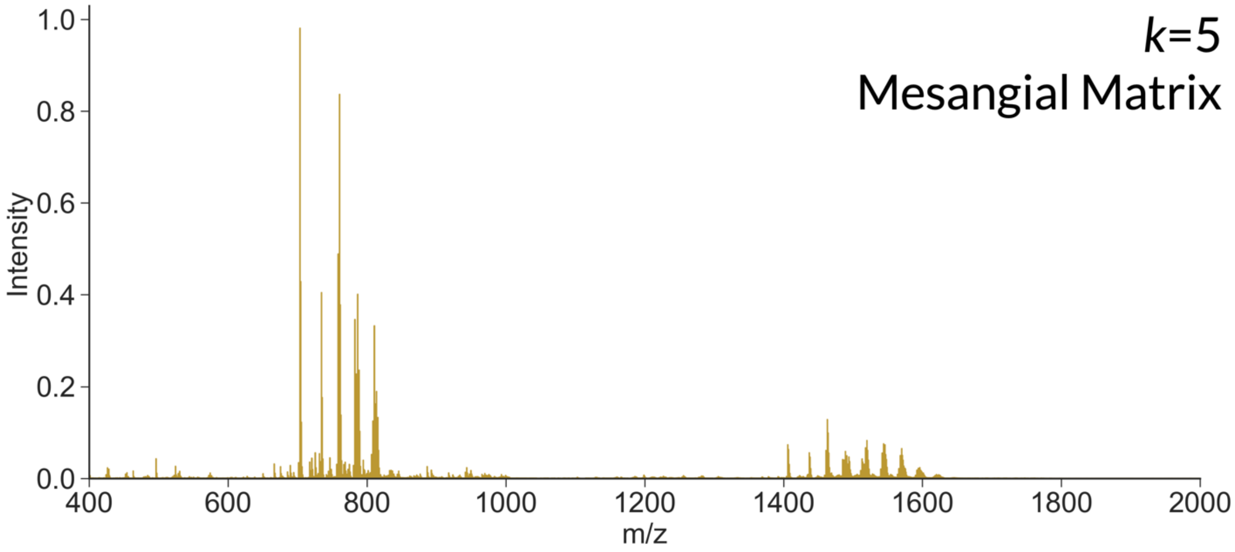


**Figure S31.** Average mass spectrum of cluster 5 (mesangial matrix-related segment).

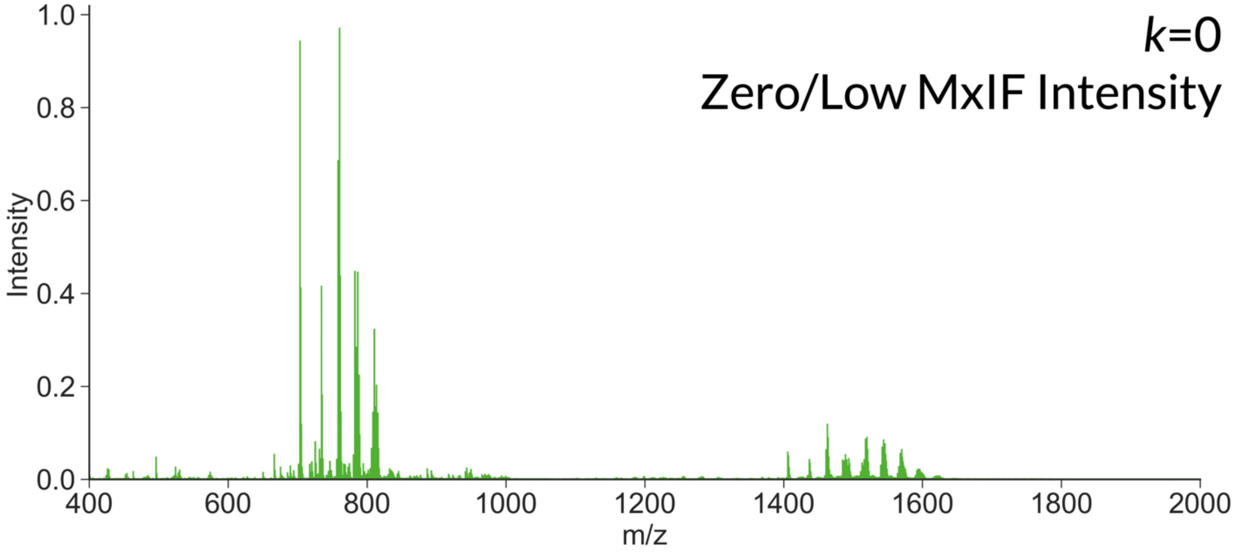


**Figure S32.** Average mass spectrum of cluster 0. This cluster is defined as regions with minimal MxIF signal as detected using the antibody marker panel described in the main paper. These low MxIF signal regions may include tissue features not stained using the included antibodies.

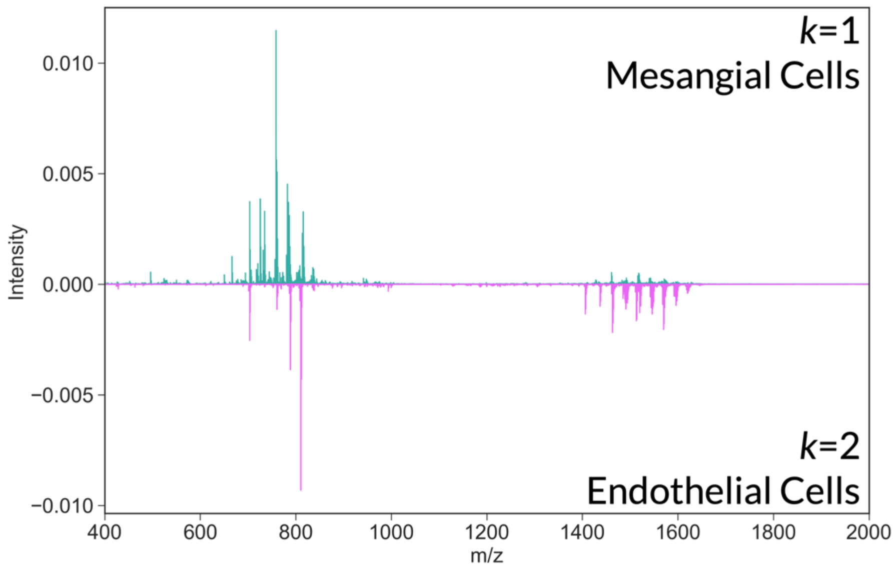


**Figure S33.** Difference spectrum showing average intensity differences between clusters 1 (mesangial cell-related segment) and 2 (endothelial cell-related segment).

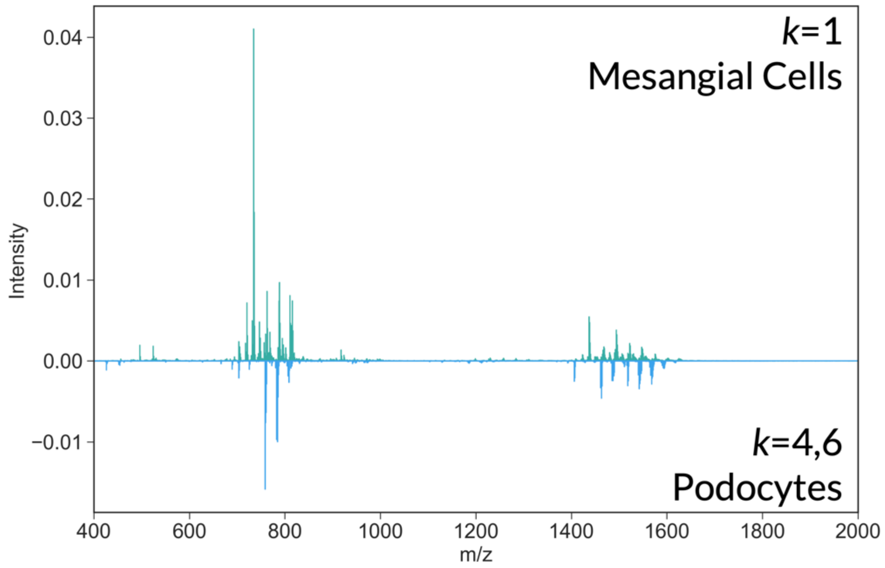


**Figure S34.** Difference spectrum showing average intensity differences between clusters 1 (mesangial cell-related segment) and a combined 4 and 6 (podocyte-related segment).

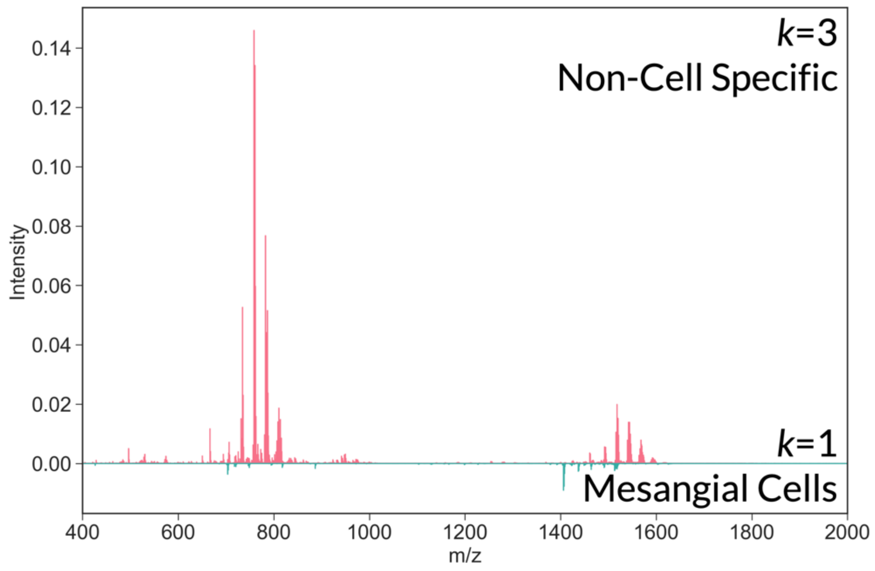


**Figure S35.** Difference spectrum showing average intensity differences between clusters 3 (non-cell specific) and 1 (mesangial cell-related segment). The non-cell specific cluster includes tissue regions with low-level signal from tensin and minimal MxIF signal detected in other channels. These regions include pixels with seemingly high background signal and low cell type specificity.

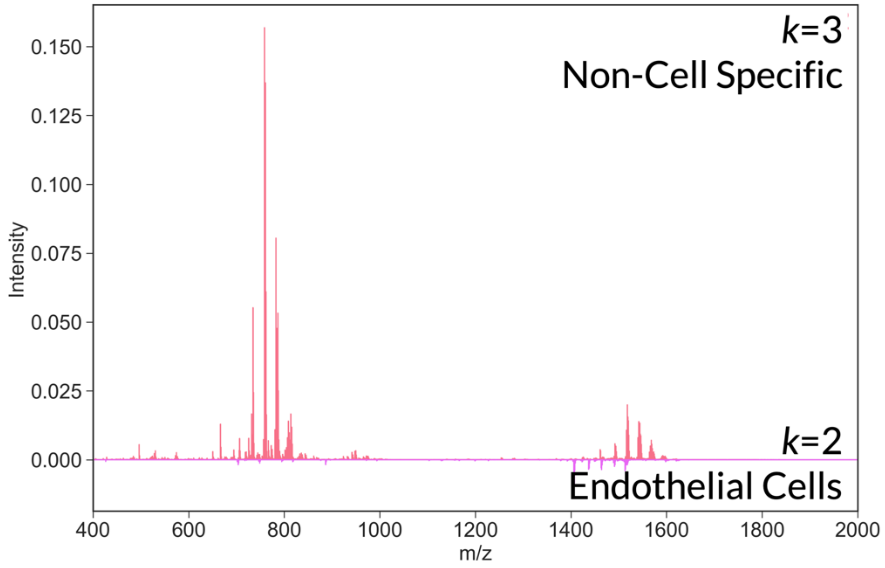


**Figure S36.** Difference spectrum showing average intensity differences between clusters 3 (non-cell specific) and 2 (endothelial cell-related segment). The non-cell specific cluster includes tissue regions with low-level signal from tensin and minimal MxIF signal detected in other channels. These regions include pixels with seemingly high background signal and low cell type specificity.

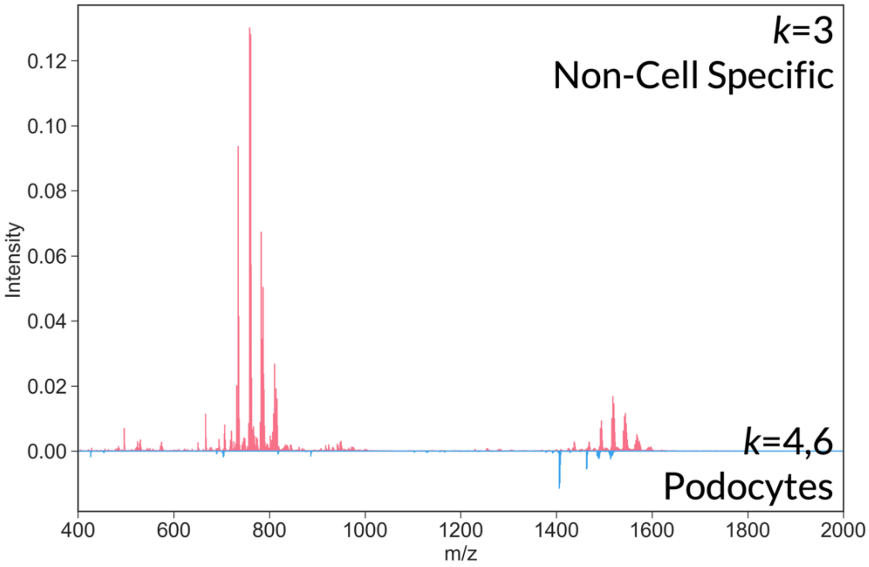


**Figure S37.** Difference spectrum showing average intensity differences between clusters 3 (non-cell specific) and a combined 4 and 6 (podocyte-related segment). The non-cell specific cluster includes tissue regions with low-level signal from tensin and minimal MxIF signal detected in other channels. These regions include pixels with seemingly high background signal and low cell type specificity.

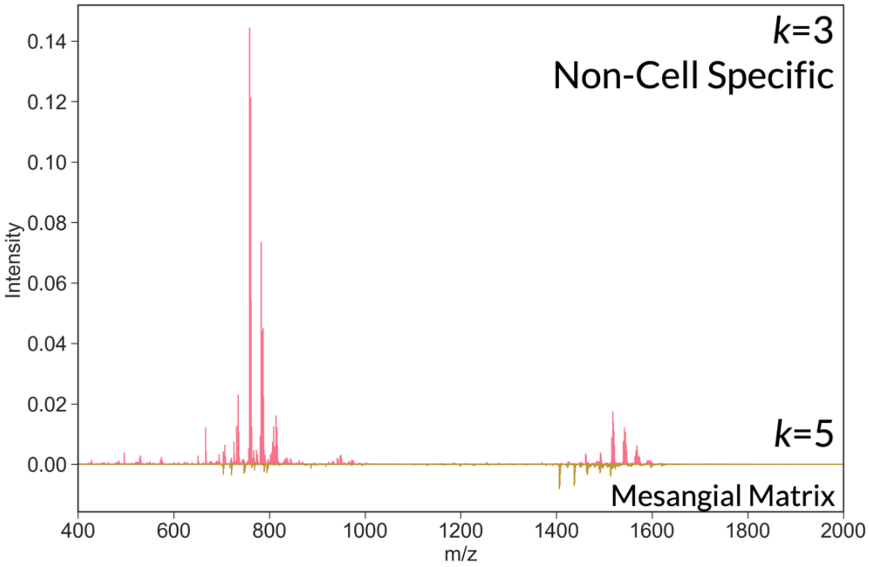


**Figure S38.** Difference spectrum showing average intensity differences between clusters 3 (non-cell specific) and 5 (mesangial matrix-related segment). The non-cell specific cluster includes tissue regions with low-level signal from tensin and minimal MxIF signal detected in other channels. These regions include pixels with seemingly high background signal and low cell type specificity.

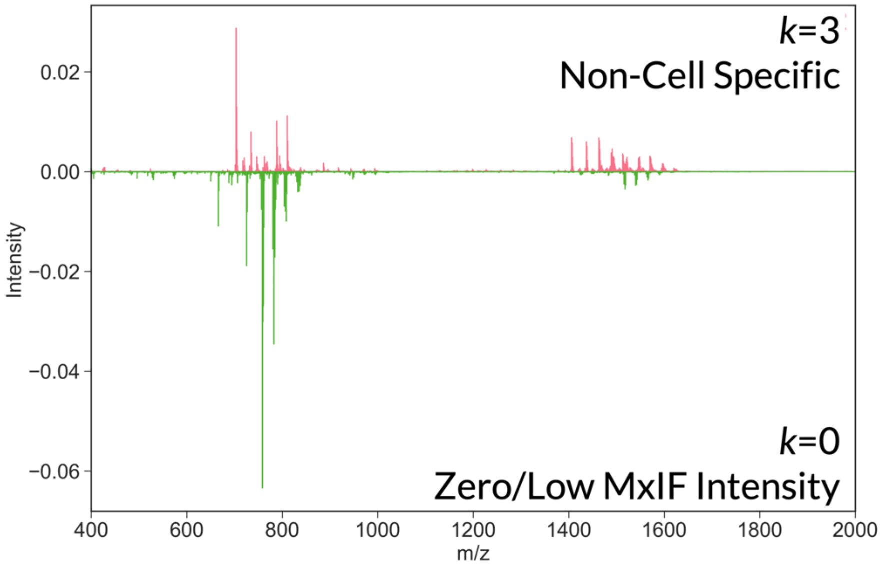


**Figure S39.** Difference spectrum showing average intensity differences between clusters 3 (non-cell specific) and 0 (zero/low MxIF intensity). The non-cell specific cluster includes tissue regions with low-level signal from tensin and minimal MxIF signal detected in other channels. These regions include pixels with seemingly high background signal and low cell type specificity. The Zero/Low MxIF intensity cluster is defined as regions with minimal MxIF signal detected using the included antibody marker panel. These low MxIF signal regions may include tissue features not stained using the included antibodies.

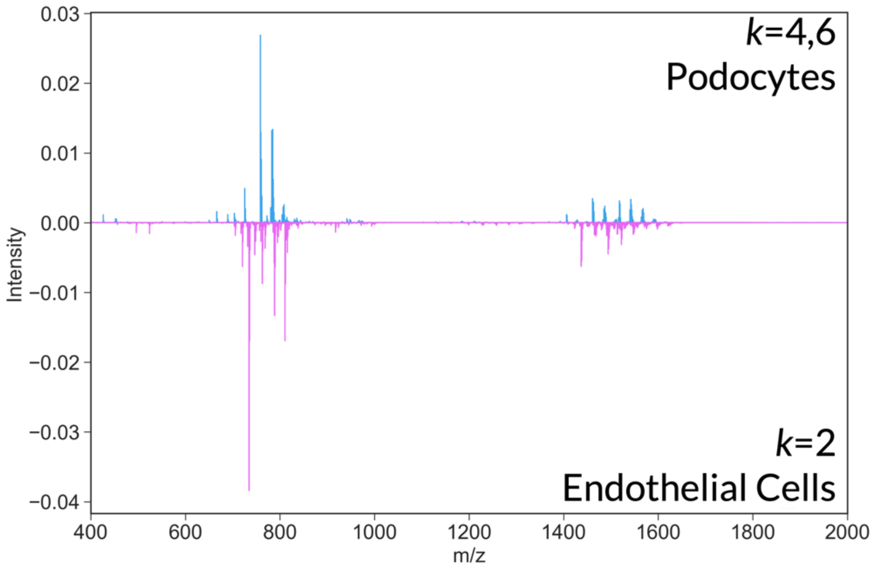


**Figure S40.** Difference spectrum showing average intensity differences between combined clusters 4 and 6 (podocyte-related segment) and 2 (endothelial cell-related segment).

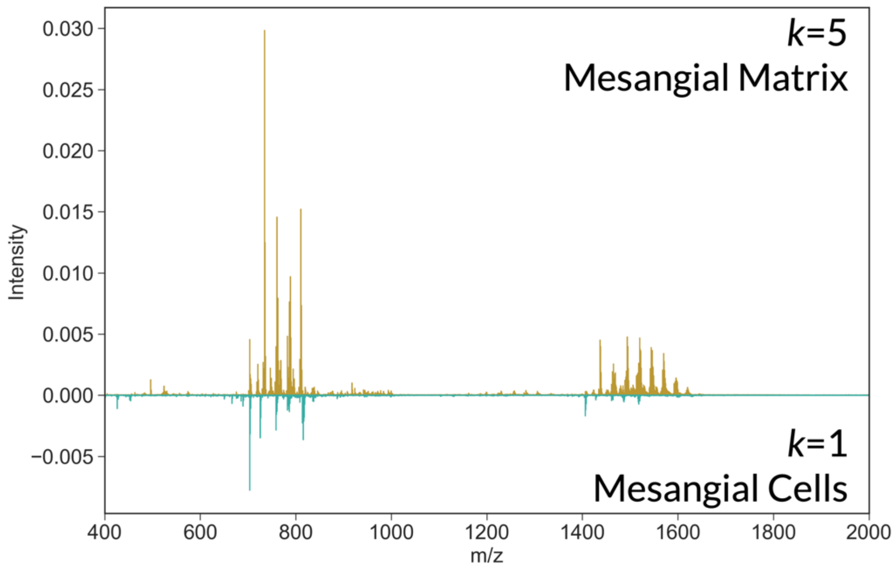


**Figure S41.** Difference spectrum showing average intensity differences between clusters 5 (mesangial matrix-related segment) and 1 (mesangial cell-related segment).

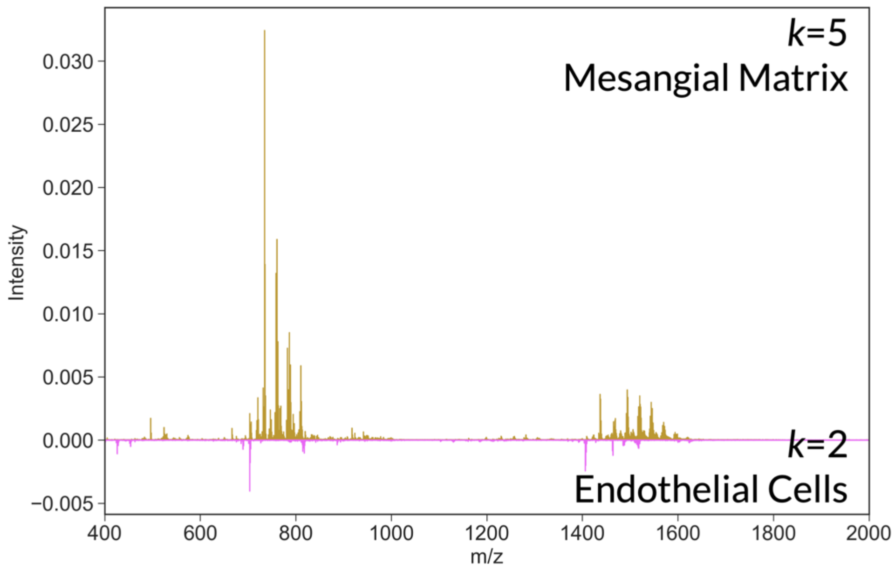


**Figure S42.** Difference spectrum showing average intensity differences between clusters 5 (mesangial matrix-related segment) and 2 (endothelial cell-related segment).

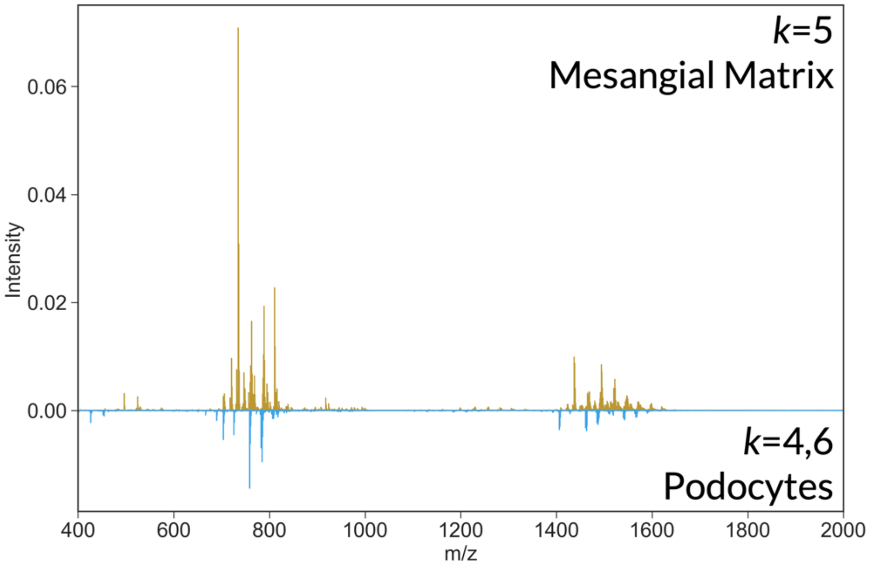


**Figure S43.** Difference spectrum showing average intensity differences between clusters 5 (mesangial matrix-related segment) and combined clusters 4 and 6 (podocyte-related segment).

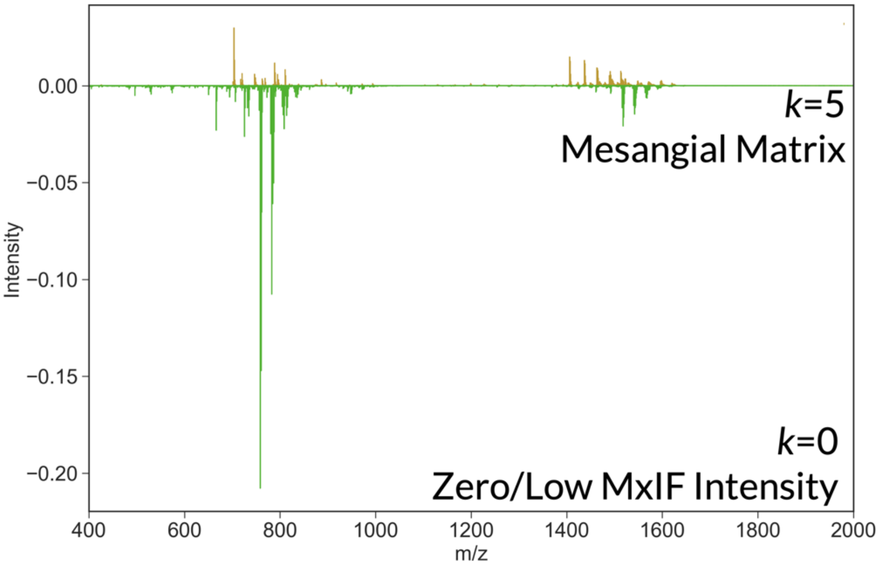


**Figure S44.** Difference spectrum showing average intensity differences between clusters 5 (mesangial matrix-related segment) and 0 (zero/low MxIF intensity). The Zero/Low MxIF intensity cluster is defined as regions with minimal MxIF signal detected using the included antibody marker panel. These low MxIF signal regions may include tissue features not stained using the included antibodies.

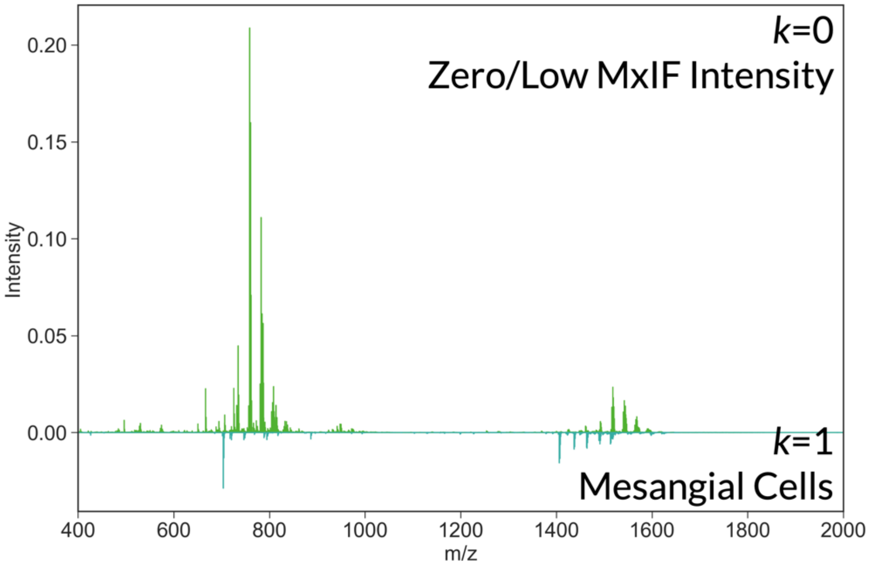


**Figure S45.** Difference spectrum showing average intensity differences between clusters 0 (zero/low MxIF intensity) and 1 (mesangial cell-related segment). The Zero/Low MxIF intensity cluster is defined as regions with minimal MxIF signal detected using the included antibody marker panel. These low MxIF signal regions may include tissue features not stained using the included antibodies.

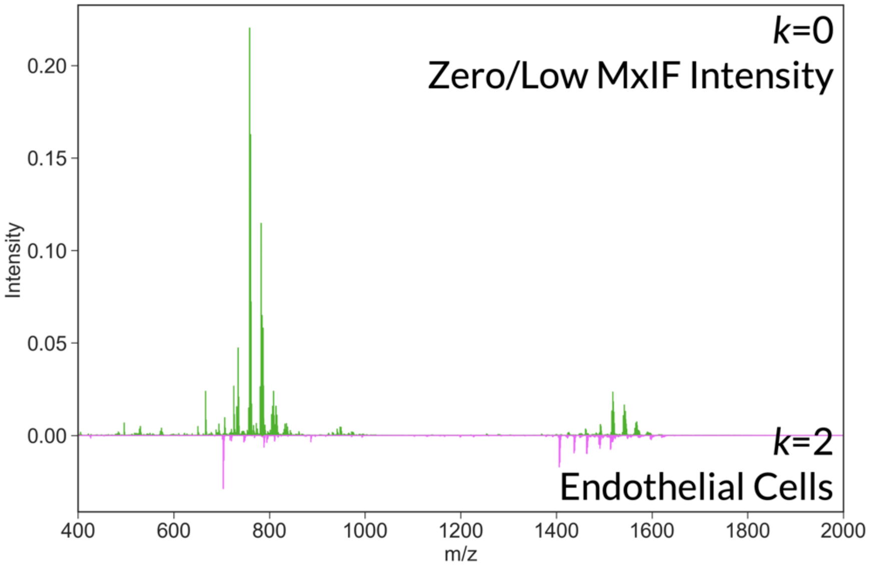


**Figure S46.** Difference spectrum showing average intensity differences between clusters 0 (zero/low MxIF intensity) and 2 (endothelial cell-related segment). The Zero/Low MxIF intensity cluster is defined as regions with minimal MxIF signal detected using the included antibody marker panel. These low MxIF signal regions may include tissue features not stained using the included antibodies.

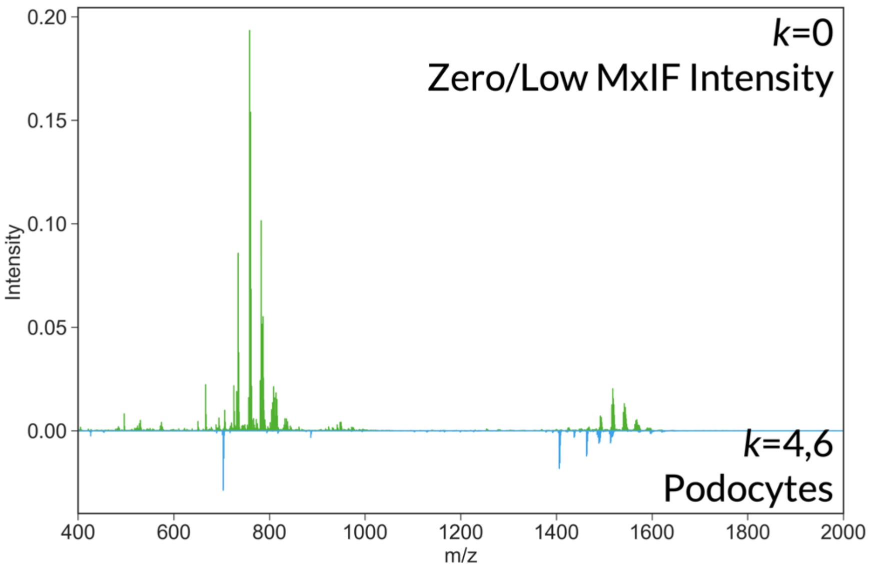


**Figure S47.** Difference spectrum showing average intensity differences between clusters 0 (zero/low MxIF intensity) and combined clusters 4 and 6 (podocyte-related segment). The Zero/Low MxIF intensity cluster is defined as regions with minimal MxIF signal detected using the included antibody marker panel. These low MxIF signal regions may include tissue features not stained using the included antibodies.

**Table S5. XGBoost Classification Model Performance Metrics per Cluster.**

| Cluster 1 - Mesangial Cells | Minimum | Maximum | Mean |
|---|---|---|---|
| Balanced Accuracy | 0.977 | 0.9783 | 0.9777 |
| F1-Score | 0.9754 | 0.9767 | 0.976 |
| Precision | 0.9973 | 0.9975 | 0.9974 |
| Recall | 0.9542 | 0.9568 | 0.9556 |
| Cluster 2 - Endothelial Cells | | | |
| Balanced Accuracy | 0.9037 | 0.9051 | 0.9042 |
| F1-Score | 0.8812 | 0.8833 | 0.8821 |
| Precision | 0.9607 | 0.9635 | 0.9625 |
| Recall | 0.813 | 0.816 | 0.8142 |
| Cluster 4 & 6 - Podocytes | | | |
| Balanced Accuracy | 0.8872 | 0.888 | 0.8876 |
| F1-Score | 0.8812 | 0.8822 | 0.8817 |
| Precision | 0.8802 | 0.8812 | 0.8806 |
| Recall | 0.8812 | 0.8841 | 0.8827 |
| Cluster 5 - Mesangial Matrix | | | |
| Balanced Accuracy | 0.9997 | 0.9999 | 0.9998 |
| F1-Score | 0.987 | 0.9996 | 0.9991 |
| Precision | 0.998 | 0.9993 | 0.9986 |
| Recall | 0.9994 | 0.9999 | 0.9996 |

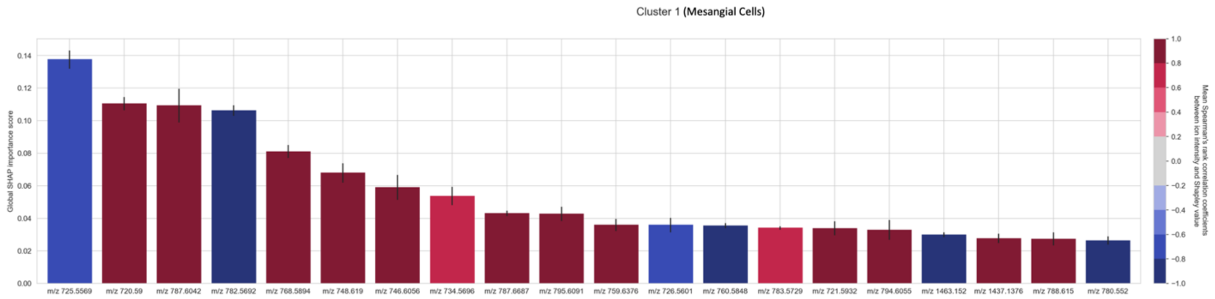


**Figure S48.** Global SHAP scores in ranked magnitude order for cluster 1 (primarily mesangial cells). The color of each bar reports whether that ion species tends to be positively (red) or negatively (blue) correlated to cluster 1.

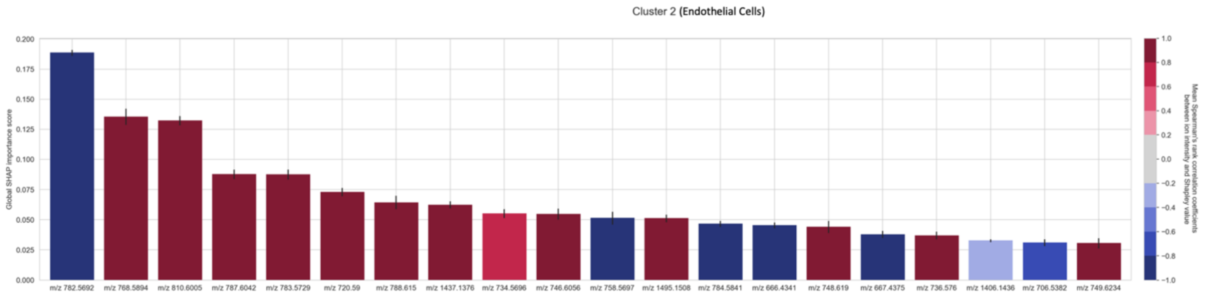


**Figure S49.** Global SHAP scores in ranked magnitude order for cluster 2 (primarily endothelial cells). The color of each bar reports whether that ion species tends to be positively (red) or negatively (blue) correlated to cluster 2.

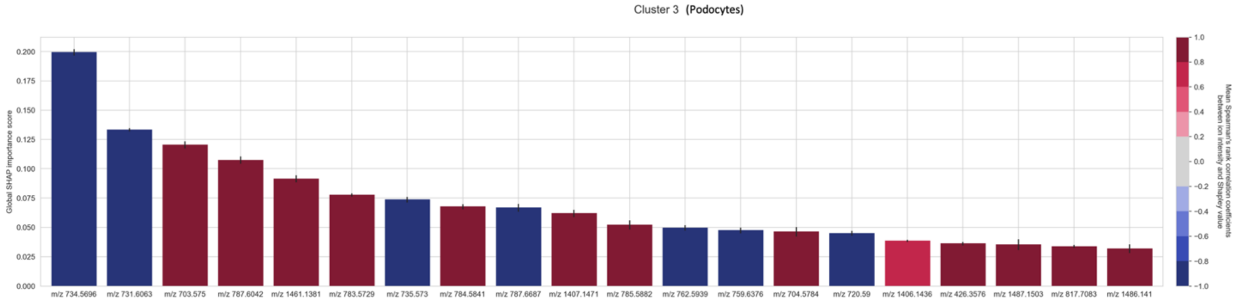


**Figure S50.** Global SHAP scores in ranked magnitude order for the combined clusters 4 and 6 (primarily podocytes). The color of each bar reports whether that ion species tends to be positively (red) or negatively (blue) correlated to clusters 4 and 6.

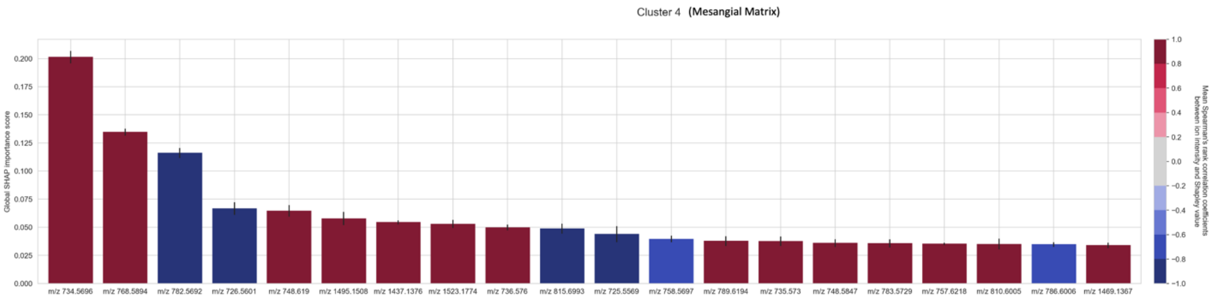


**Figure S51.** Global SHAP scores in ranked magnitude order for cluster 5 (primarily mesangial matrix). The color of each bar reports whether that ion species tends to be positively (red) or negatively (blue) correlated to cluster 5.

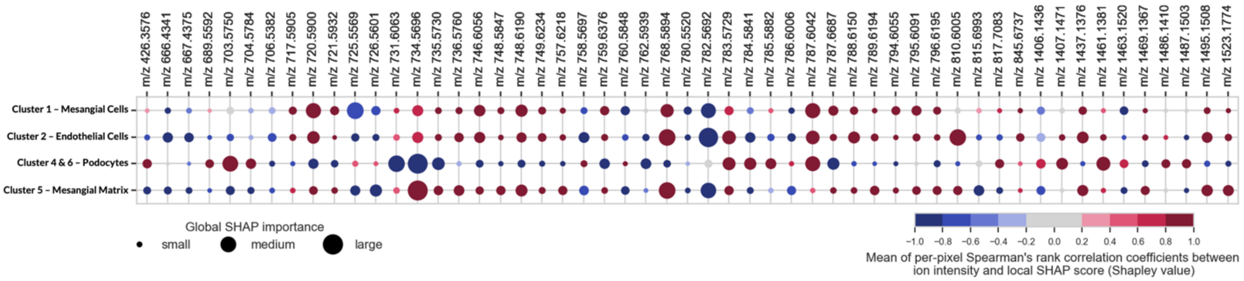


**Figure S52.** Full SHAP bubble plot, with m/z values and including isotopes and lipid dimers. The bubble plot summarizes the biomarker candidates for the glomerular segmentations and their dominant cell types. The size of each bubble indicates the global SHAP importance of a given ion species (column) to recognizing a given glomerular subarea (and its dominant cell type) (row), and the color indicates a positive (red) or negative (blue) correlation of the ion species abundance to that cluster's recognition. From this analysis, it is shown that every glomerular segmentation has its own unique profile of relevant ion species.

**Table S6.** Identifications of SHAP *m/z* values

| MALDI m/z | LC-MS/MS m/z | Theoretical m/z | MALDI ppm Error | LC-MS/MS ppm Error | Identification | Adduct Type |
|---|---|---|---|---|---|---|
| 426.3576 | 426.3568 | 426.3578 | -0.5 | -2.3 | CAR (18:1) | $[M+H]^+$ |
| 666.4341 | 666.4305 | 666.4341 | 0.0 | -5.4 | PS (27:0) | $[M+H]^+$ |
| 667.4375 | x | x | x | x | Isotope | x |
| 689.5592 | 689.5605 | 689.5592 | 0.0 | 1.9 | CerPE (d36:1) | $[M+H]^+$ |
| 703.575 | 703.5757 | 703.5748 | 0.3 | 1.3 | SM (d34:1) | $[M+H]^+$ |
| 704.5784 | x | x | x | x | Isotope | x |
| 706.5382 | 706.5381 | 706.5381 | 0.1 | 0.0 | PC (30:0) | $[M+H]^+$ |
| 717.5905 | x | 717.5905 | 0.0 | x | *CerPE (38:1) | $[M+H]^+$ |
| 720.59 | x | 720.5902 | -0.3 | x | *PC (O-32:0) | $[M+H]^+$ |
| 721.5932 | x | x | x | x | Isotope | x |
| 725.5569 | x | 725.556796 | 0.1 | x | *SM (d34:1) | $[M+Na]^+$ |
| 726.5601 | x | x | x | x | Isotope | x |
| 731.6063 | 731.6072 | 731.6061 | 0.3 | 1.5 | SM (d36:1) | $[M+H]^+$ |
| 734.5696 | 734.5672 | 734.5694 | 0.3 | -3.0 | PC (32:0) | $[M+H]^+$ |
| 735.573 | x | x | x | x | Isotope | x |
| 736.576 | x | x | x | x | Isotope | x |
| 746.6056 | 746.603 | 746.6058 | -0.3 | -3.8 | PC (O-34:1) | $[M+H]^+$ |
| 748.5847 | 748.5851 | 748.5851 | -0.5 | 0.0 | PC (33:0) | $[M+H]^+$ |
| 748.619 | 748.6198 | 748.6215 | -3.3 | -2.3 | PC (O-34:0) | $[M+H]^+$ |
| 749.6234 | x | x | x | x | Isotope | x |
| 757.6218 | 757.623 | 757.6218 | 0.0 | 1.6 | SM (d38:2) | $[M+H]^+$ |
| 758.5697 | 758.5699 | 758.5694 | 0.4 | 0.7 | PC (34:2) | $[M+H]^+$ |
| 759.6376 | 759.6359 | 759.6374 | 0.3 | -2.0 | SM (d38:1) | $[M+H]^+$ |
| 760.5848 | 760.5842 | 760.5851 | -0.4 | -1.2 | PC (34:1) | $[M+H]^+$ |
| 762.5939 | x | x | x | x | Isotope | x |
| 768.5894 | x | 768.5902 | -1.0 | x | *PC (O-36:4) | $[M+H]^+$ |
| 780.552 | 780.5558 | 780.5538 | -2.3 | 2.6 | PC (36:5) | $[M+H]^+$ |
| 782.5692 | 782.5687 | 782.5694 | -0.3 | -0.9 | PC (36:4) | $[M+H]^+$ |
| 783.5729 | x | x | x | x | Isotope | x |
| 784.5841 | 784.5872 | 784.5851 | -1.3 | 2.7 | PC (36:3) | $[M+H]^+$ |
| 785.5882 | x | x | x | x | Isotope | x |
| 786.6006 | 786.5972 | 786.6007 | -0.1 | -4.4 | PC (36:2) | $[M+H]^+$ |
| 787.6042 | x | x | x | x | Isotope | x |
| 787.6687 | 787.6688 | 787.6687 | 0.0 | 0.1 | SM (d40:1) | $[M+H]^+$ |
| 788.615 | 788.618 | 788.6164 | -1.8 | 2.0 | PC (36:1) | $[M+H]^+$ |
| 789.6194 | x | x | x | x | Isotope | x |
| 794.6055 | x | 794.6058 | -0.4 | x | *PC (O-38:5) | $[M+H]^+$ |
| 795.6091 | x | x | x | x | Isotope | x |
| 796.6195 | 796.6172 | 796.6215 | -2.5 | -5.4 | PC (O-38:4) | $[M+H]^+$ |
| 810.6005 | x | 810.6007 | -0.2 | x | *PC (38:4) | $[M+H]^+$ |
| 815.6993 | 815.6976 | 815.7001 | -1.0 | -3.1 | SM (d42:1) | $[M+H]^+$ |
| 817.7083 | x | x | x | x | Unknown | x |
| 845.6737 | x | x | x | x | Unknown | x |
| 1406.1436 | x | x | x | x | Lipid Dimer | x |
| 1407.1471 | x | x | x | x | Lipid Dimer | x |
| 1437.1376 | x | x | x | x | Lipid Dimer | x |
| 1461.1381 | x | x | x | x | Lipid Dimer | x |
| 1463.152 | x | x | x | x | Lipid Dimer | x |
| 1469.1367 | x | x | x | x | Lipid Dimer | x |
| 1486.141 | x | x | x | x | Lipid Dimer | x |
| 1487.1503 | x | x | x | x | Lipid Dimer | x |
| 1495.1508 | x | x | x | x | Lipid Dimer | x |
| 1523.1774 | x | x | x | x | Lipid Dimer | x |

*****Identified with exact mass from MALDI *m/z*
